# Targeting the RNA regulator HuR in acute traumatic brain injury mitigates secondary injury and improves functional outcome by suppressing neuroinflammation

**DOI:** 10.64898/2026.10.22.761057

**Authors:** Mohammad Amir Husain, Ben Sheline, Reed Smith, Oanh T. P. Trinh, Ying Si, Annye P. Bennett, Mark Bolding, Shreyas Keshava, L. Burt Nabors, Natalia Filippova, Peter H. King

## Abstract

Traumatic brain injury (TBI) affects ∼2.5 million people annually in the US. Minutes after TBI, microglia and astrocytes initiate a neuroinflammatory cascade which promotes secondary injury via cytotoxicity, blood-brain barrier damage, edema, and immune cell recruitment. HuR, an RNA regulator, plays a critical role in the glial response by promoting the expression of proinflammatory mediators including IL-6, IL-1β, TNF-α, COX-2, CCL2, and CXCL1 via adenine and uridine-rich elements in the 3’ untranslated region of their mRNAs. Here, we assessed SRI-42127, a small molecule HuR inhibitor developed by our group, on neuroinflammatory responses, secondary injury, and functional outcome after TBI. Male mice underwent controlled cortical impact (CCI) followed by peripheral administration of SRI-42127. Functional outcomes assessed included: motor function (balance beam/rotarod) and memory function (object location/recognition). Edema, lesion volume, and BBB permeability were assessed by MRI, brain water content, and Evans blue dye extravasation. HuR nucleocytoplasmic translocation, neuronal loss/apoptosis, microglial and astrocyte activation, and neutrophil infiltration were quantified by immunohistochemistry. Proinflammatory mediator induction was assessed by RT-qPCR and/or immunoblot. Results showed that SRI-42127 treatment improved spatial memory and fine motor performance in the first week in parallel with a reduction in edema, vascular permeability, lesion size, and neuronal loss/apoptosis. There was attenuated microglial activation and astrogliosis, decreased neutrophil infiltration, and suppression of proinflammatory mediator induction. Mechanistically, SRI-42127 inhibited HuR nucleocytoplasmic translocation in microglia and astrocytes. In summary, SRI-42127 potently suppresses neuroinflammation and secondary injury after TBI, improves functional outcomes, and identifies HuR as a new therapeutic target in TBI.

## 1. Introduction

Traumatic brain injury (TBI) affects 2.5 million people in the US annually, and over 5 million experience long-term sequelae including neurological and behavioral impairment, epilepsy, and an elevated risk of neurodegenerative disease (Barker et al., 2023; Simon et al., 2017). In TBI, the primary injury produces immediate impairment through physical impact, but this is rapidly followed by secondary tissue injury stemming from separate pathophysiological processes including blood-brain barrier (BBB) disruption, vascular permeability/edema and ischemia (Jassam et al., 2017). Secondary injury can progress over months, and even longer in a subset of patients. Convergence of these processes often leads to worse tissue damage than the primary injury (Wu et al., 2026).

A major driver of secondary injury stems from a toxic cascade of neuroinflammation which is triggered within minutes of TBI and peaks at 24-72 h (Jassam et al., 2017; Simon et al., 2017). The breadth of proinflammatory mediators released in this phase is quite large and includes TNF-α, IL-1β, IL-6, nitric oxide (iNOS), PGE2 (COX-2), and MMPs. Glial and neuronal signaling activates NLRP3 which further amplifies tissue injury by caspase-1 and 3—mediated apoptosis (Irrera et al., 2020). Chemokines released by these cells, including CXCL1 and CCL2, drive recruitment of neutrophils and monocytes via a damaged BBB (Semple et al., 2010). Neutrophils are the earliest infiltrating immune cells, peaking at 24 h post injury, and promoting secondary tissue injury via extracellular trap formation (Gyoneva and Ransohoff, 2015; Slough et al., 2025; Vaibhav et al., 2020). Reactive oxygen species (ROS) and reactive nitrogen species (RNS) are cytotoxic byproducts of neuroinflammation and are produced by many cell types, ranging from microglia, astrocytes, and endothelial cells in the central nervous system (CNS) to infiltrating neutrophils and monocyte/macrophages from the periphery (Fesharaki-Zadeh, 2022). Not surprisingly, therapeutic targeting of the immune response in TBI has failed due to the many overlapping effects of these mediators and their secretion by multiple cell types (Jassam et al., 2017).

Despite the diversity of proinflammatory mediators induced in TBI, a common node for their regulation is at the mRNA level where specific adenine and uridine-rich elements (ARE) in the 3’ untranslated region (UTR) govern expression through an interaction with cellular ARE binding proteins (AUBP) (Guha et al., 2023; Guha et al., 2022). HuR is one of these AUBPs and is a major positive regulator of proinflammatory mediators across multiple cell types (Guha et al., 2025; Guha et al., 2022; Majunder et al., 2022; Wang et al., 2019). In prior work, we have extensively characterized HuR in microglia and astrocytes and have found that it positively regulates a broad number of proinflammatory mediators linked to acute TBI (Chellappan et al., 2022; Kwan et al., 2017a; Matsye et al., 2017; Wang et al., 2019). Genetic silencing or chemical inhibition of HuR in microglia attenuates chemoattraction of neutrophils and monocytes to the central nervous system (CNS) in a lipopolysaccharide (LPS)-induced model of neuroinflammation (Chellappan et al., 2022; Matsye et al., 2017). We have shown that transgenic overexpression of HuR in astrocytes increases vascular permeability and exacerbates secondary tissue damage after spinal cord injury (SCI) and stroke (Ardelt et al., 2017; Kwan et al., 2017a; Kwan et al., 2017b). Given that these phenotypes are major drivers of secondary injury and functional loss in TBI, including long term sequelae of neurodegeneration (Vazquez-Rosa et al., 2020), we hypothesized that inhibition of HuR may be a viable therapeutic strategy in TBI. Our group has developed a small molecule, SRI-42127, that inhibits two key functional properties of HuR necessary for its activity: homodimerization and cytoplasmic translocation (Filippova et al., 2021; Filippova et al., 2017; Meisner et al., 2007). We previously validated this mechanism of action in activated microglia and astrocytes where SRI-42127 selectively blocked HuR nucleocytoplasmic translocation and potently suppressed production of proinflammatory mediators (Chellappan et al., 2022). Its potential as a therapeutic target in acute CNS injury was recently demonstrated by our group where we showed that SRI-42127 blocked the induction of proinflammatory mediators in a model of SCI and significantly mitigated loss of motor function and neuropathic pain (Husain et al., 2025). The current study aimed to investigate the effect of HuR inhibition with SRI on neuroinflammatory responses in acute TBI and the impact on functional outcome.

## 2. Methods

### 2.1. Animals and CCI model of traumatic brain injury

All animal procedures were reviewed and approved by the Institutional Animal Care and Use Committee (IACUC) at the University of Alabama at Birmingham (UAB; Protocol Number-23116). These experiments were performed in strict accordance with the National Institutes of Health (NIH) Guide for the Care and Use of Laboratory Animals. The study design, animal monitoring, and data reporting were conducted in alignment with the ARRIVE guideline. The CCI model was chosen for this study because of its reproducibility, precise control over injury parameters, and ability to target specific brain regions such as the cortex and hippocampus (Osier and Dixon, 2016). Adult male mice C57BL/6 (10–14 weeks) underwent CCI (RWD 68099 II) under isoflurane anesthesia using a stereotaxic frame. A 4-mm craniotomy was performed over the right parietal cortex (2 mm lateral, 2 mm posterior to bregma) targeting the dorsal hippocampus and posterior sensorimotor cortex to assess cognitive and motor deficits. Injury was induced with a 3-mm impactor (3.5 m/s velocity, 2.0 mm depth, 200 ms dwell time), while sham controls received craniotomy only. Intraperitoneal (i.p.) administration of SRI-42127 (10 mg/kg) or vehicle was initiated 1 h post-injury. One dose of subcutaneous buprenorphine was administered prior to injury. For molecular studies, SRI-42127 (10 mg/kg) and vehicle were administered i.p. every 2 h beginning 1 h after injury for 4 doses. The cohort for behavioral studies received 4 doses daily every 6 h, starting 1 h after injury, for 5 days similar to our prior study with SCI (Husain et al., 2025). Sham mice (craniotomy without impact; vehicle-treated) were included as controls for the balance beam experiments and memory testing, and pre-injury baselines served as internal controls for all other behavioral tests. The contralateral hemisphere were used as internal controls for the RNA and histological analyses. The contralateral hemisphere was validated as a non-injured reference by comparison with sham controls in the qPCR experiments (Suppl. Fig. 8).

### 2.2. RNA isolation, qPCR and Western blot

The brain was harvested with a 4-mm tissue puncture at the lesion site, with a corresponding sample collected from the contralateral side as a control. RNA was isolated using TRIzol reagent as per manufacturer’s protocol (Thermo Fisher Scientific). RNA was precipitated using isopropanol (Sigma) and concentrations were measured by Nanodrop (Thermo Scientific, Waltham, MA). Complementary DNA was reverse transcribed from 2 μg of total RNA using a reverse transcription kit (Thermo Fisher Scientific). Quantitative Real Time PCR (RT-qPCR) was conducted using the ViiA7-Real-Time PCR system as previously described (Chellappan et al., 2022). For western blot analysis, whole cell lysates were prepared using T-PER (Thermofisher), and protein quantification was conducted with a BCA protein assay kit (Thermofisher). Samples (80 µg) were electrophoresed in a 4%–20% mini-PROTEAN gel (Biorad) and transferred to a nitrocellulose membrane. Blots were immunostained with a COX-2 antibody (Santa Cruz Biotechnology, USA Inc) at 1:2000 and a GAPDH antibody (14C10, Cell Signaling) at 1:3000. Band intensities were quantified from three biological replicates (n = 3 mice per group) using densitometry by Image Lab 6.0.1 (Bio-Rad Laboratories, USA Inc.). With the Volume Tools, a rectangle of identical size was drawn around each band. COX-2 signal was normalized to GAPDH in the same lane and expressed relative to the contralateral side of vehicle-treated mice, set at 1.0.

### 2.3. Immunohistochemistry

Mice were euthanized at 24 h and perfused with an ice-cold phosphate buffer saline (PBS) solution for 2 min followed by 4% paraformaldehyde (PFA) solution for 5 min (Kwan et al., 2017a). The entire brain was then extracted and post-fixed in PFA for 24 h to ensure complete tissue fixation. The fixed brain was then incubated in a PBS solution with 10% sucrose for cryoprotection and transferred to a 30% sucrose in PBS solution for 48 h. Thereafter, the brain was removed from the sucrose solution and cryo-molded in OCT. A cryostat was used to cut samples into 15 μm coronal serial sections.

Sections were washed with PBS and blocked in 10% goat serum for 1 h. Samples were incubated overnight with primary antibodies (GFAP, 1:500, Z0334, Dako, Carpinteria, CA), (Iba1, 1:500, FUJIFILM Wako Chemicals U.S.A. Corporation), (NeuN, 1:400, ABN78, Sigma-Aldrich, USA Ltd.), (HuR, 1:500, sc-5261, Santa Cruz Biotechnology, USA Inc), (Cleaved-Caspase 3, 1:400, 68773, Proteintech), (Elastase, 1:200, E8U3X, Cell Signalling), (MPO, 1:200, AB9535, Abcam). The following day, slide sections were washed and incubated with Alexa fluorophore-conjugated secondary antibodies (Goat anti mouse, A28175, Thermo Fisher Scientific and Goat anti rabbit Cy3, 111-165-144, Jackson Immuno Research) for 2 h, mounted with antifade DAPI + mountant (ThermoFisher Scientific P36966).

Quantitative assessment of HuR localization was done using our previously published method (Chellappan et al., 2022). Fiji software was utilized for image analysis, and a minimal threshold was established in order to quantify the mean fluorescence intensity (FI). FI was quantified and normalized with the DAPI FI for the same region and expressed as arbitrary units (AU). All immunohistochemical analyses were performed by an investigator blinded to the experimental groups. Quantification of HuR localization, GFAP, Iba1, and NeuN was performed using Fiji as described above. Neuronal apoptosis was measured as the percentage of NeuN⁺ neurons co-labeled with cleaved caspase-3 in the perilesional cortex, calculated as (NeuN⁺/cleaved caspase-3⁺ cells /total NeuN⁺ cells) × 100. Cleaved caspase-3-positive neurons and MPO– and neutrophil elastase-positive cells within the perilesional area were manually quantified from 60× images by an observer blinded to the experimental conditions. Data from four biological replicates per group (n = 4) were used for statistical analysis.

### 2.4. Analysis of microglial and astrocyte morphology

Morphological assessments were based on methods previously described (Husain et al., 2025). Briefly, Fiji software (https://imagej.net/) was used to convert all 60x photomicrographs of brain sections to binary and skeletonized images and were processed using AnalyzeSkeleton plugin (https://imagej.net/plugins/analyze-skeleton/). To analyze the number of branches and the total branch length per microglia or astrocyte, brightness and contrast were changed by updating the Fiji lookup table while leaving pixel values unchanged. This is followed by a despeckle step to remove salt-and-pepper noise. Photomicrographs were then converted to binary using threshold tool followed by skeletonizing the image using the toolbar Skeletonize and AnalyzeSkeleton to collect data on the branch length. The total branch length per microglia and astrocyte was calculated by multiplying the total number of branches by the average branch length. Data were obtained from 20 microglial cells from vehicle control (n=4) and 21 microglial cells from SRI-42127 (n=4), and from 10 astrocyte cells from vehicle control (n=4) and 11 astrocyte cells from SRI-42127 (n=4). The soma area was calculated using the Fiji multipoint area selection tool.

### 2.5. Histological assessment of tissue injury and Edema

Nissl staining and MRI images were employed to evaluate the cortical lesion area. A series of sections were stained with cresyl violet to delineate the spared cortex. The lesion area of the cavity was subsequently calculated using Fiji software. MRI was performed on a 9.4T Bruker BioSpec 94/20 scanner (ParaVision 360 v3.6) using a volume resonator for transmission and a receive-only surface array coil. T2-weighted images (T2-WI) were acquired with a 2D RARE sequence (TR 2500 ms, effective TE 40 ms, RARE factor 8, 6 averages) with 23 coronal slices of 0.5 mm thickness, a 25 × 25 mm field of view and a 256 × 256 matrix (98 µm in-plane resolution), for a scan time of 8 minutes. For T2 scans, the hyperintense area was identified as the lesion area and manually outlined using Fiji. The lesion volume was calculated using the Ratios of Ipsilateral and Contralateral Cerebral Hemispheres (RICH) method, as described earlier (Boyko et al., 2013), with the following formula: Corrected lesion volume = Lesion volume × Contralateral hemisphere size / Ipsilateral hemisphere size. MRI scans were utilized to measure edema post-TBI using T2WI sequences. Additionally, the dry/wet method was employed to analyze brain edema. The brain was harvested, and since edema is localized around the impact site, a 4-mm tissue punch was carefully extracted from the injured hemisphere and a mirror punch from the contralateral hemisphere, as previously described (Su et al., 2017). Dry weight was obtained after drying in an oven at 95°C for 24 h. The percentage of tissue water content was calculated as follows: [(wet weight – dry weight) / wet weight] × 100%. Nissl-stained sections and T2-weighted MRI images were analyzed by an investigator blinded to the experimental groups.

### 2.6. BBB disruption analysis

BBB damage was assessed by quantifying Evans blue (EB) dye uptake, as described by (Xiang et al., 2026; Zhang et al., 2022). At 24 h after TBI, a 2% Evans blue dye solution was administered intravenously via the retro-orbital venous sinus at a dosage of 4 mL/kg at 2 h prior to euthanasia (Steel et al., 2008). The mice were anesthetized and perfused with pre-cooled PBS. Tissue samples, measuring 4 mm, were collected from both the injured and corresponding contralateral hemispheres. The brain tissue was sectioned and placed in a tube containing 300 µL of dimethylformamide (Sigma-Aldrich). All samples were incubated at 60° C for 48 h and subsequently centrifuged at 14,000 rpm for 30 min to obtain the supernatants. Standard solutions of Evans blue were prepared, and the absorbance of both the sample supernatants and the standard solutions were measured at 620 nm using a microplate reader (BioTek). Evans blue dye content in each sample was determined using the standard curve, normalized to the brain weight, and expressed as μg/g tissue. The contralateral side served as an internal control for each mouse.

### 2.7. Novel object recognition (NOR) and object location test (OLT)

Open field test was performed to measure NOR and OLT. The experimental chamber consists of a Plexiglas box with dimensions (16 in. x 16 in x 12 in). The experiment was monitored by an automated activity EthoVision XT video monitoring system. Both NOR and OLT are spontaneous exploration tasks for mice (Denninger et al., 2018). NOR examines object recognition memory function and OLT is a dorsal hippocampus behavioral assay to study spatial memory. Mice were habituated to the empty arena before surgery and again immediately before familiarization. Training (familiarization) was performed 24 h after TBI where mice explored two identical objects for 10 min. Memory was then tested at two retention intervals after training: 90 min following training, (short-term memory) or 48 h (long-term memory). At testing, one object was replaced with a novel object (NOR) or moved to a new location (OLT), and the percentage of exploration time per object and the discrimination index were recorded. (Antunes and Biala, 2012; Cohen and Stackman, 2015). Sham mice (n = 4) were tested in the short-term OLT.

### 2.8. Balance Beam

The apparatus consists of a round horizontal beam of 100 cm long and 1.2 cm in diameter and elevated 50 cm above the ground. A hollow black escape box was attached to one end of the beam. One fluorescent lamp (60W) was used to illuminate the beam from the start side. One week prior to CCI, mice were thoroughly trained to traverse the beam from start point to end point (black box). Baseline measurements were taken one day before the CCI. On the trial day after CCI mice were placed at the start point and allowed to walk for 1 min for 2 trials. The test was recorded by video camera. We measured the time to cross the beam from one end to other and the number of foot slips at days 1, 3, 5 and 7 after TBI. A mean value was calculated from the two trials for each mouse. Sham mice (n = 4) were tested at baseline and 1 day after surgery.

### 2.9. Rotarod

Mice were trained one week before CCI by placing them on the revolving rotor (4 rpm) and allowing them to acclimate for 2 min. Baseline measures were acquired one day prior to CCI. On the testing day, the rod accelerated from 4 to 60 rpm over 2 min and the latency to fall off was measured in 3 trials on each testing day, allowing them 5 min of rest between each trial. The average time for the 3 trials was then used for statistical analysis to calculate a single score for each mouse at days 1, 3, 5, and 7 after TBI. Behavioral analyses (NOR, OLT, balance beam and rotarod) were performed by an observer blinded to treatment group.

### 2.10. Statistics

All statistical analyses were performed by Graphpad (GraphPad Software Inc.). The mean values are shown on graphs along with error bars. Error bars indicate the standard deviation and standard error as indicated. An unpaired two-tailed Welch’s t test was applied to analyze the difference between two groups, whereas two-way ANOVA with Tukey post hoc test was applied to compare multiple groups. A p-value less than 0.05 was considered statistically significant. To identify outliers, Grubbs test was applied with a significance threshold of α = 0.05.

## 3. Results

### 3.1. SRI-42127 treatment improves spatial memory and fine motor recovery after CCI

Mice subjected to CCI were treated with SRI-42127 (10mg/kg i.p.) every 6 h for 5 days starting 1 h post-injury **(Fig. 1A)**. Cognitive function was assessed using open field-based object recognition and object location tests. Briefly, mice explore two identical objects in an open field for 10 min (recorded via Ethovision XT). On the retention day, one object is replaced (NOR) or relocated (OLT) to assess recognition or spatial memory, respectively, at 90 min (short-term) or 48 h (long-term) post-training (Antunes and Biala, 2012; Cohen and Stackman, 2015). In the object location test, representative heat maps of exploration during familiarization and testing phases for short-term memory show that SRI-42127-treated (D) mice display a clear preference for the moved object (white arrowheads) at a novel location, while vehicle (V) mice show no location preference (**Fig. 1B**). Consistent with this, vehicle-treated TBI mice failed to show preference for the moved object versus the familiar object, as measured by % time, in both short– and long-term memory tests (**Fig. 1C**). Mice also demonstrated a negative discrimination index indicating impaired spatial memory consistent with dorsal hippocampal dysfunction (**Fig. 1D**) (Paterno et al., 2018). In contrast, SRI-42127 significantly increased the time spent exploring the moved object (p < 0.0001) and had a positive discrimination index, suggesting restoration of hippocampal-dependent spatial memory **(Fig, 1 C,D).** In contrast, sham mice showed a clear preference for the moved object (P = 0.0058) and a positive discrimination index (Suppl. Fig. 1). In the object recognition test, both vehicle– and SRI-42127-treated mice retained recognition memory, consistent with preserved cortical-hippocampal and prefrontal network, specifically the perirhinal and medial prefrontal cortices, which collectively mediate non-spatial object identity recognition (**Fig. 1 E,F**) (Broadbent et al., 2010).

**Fig. 1:**
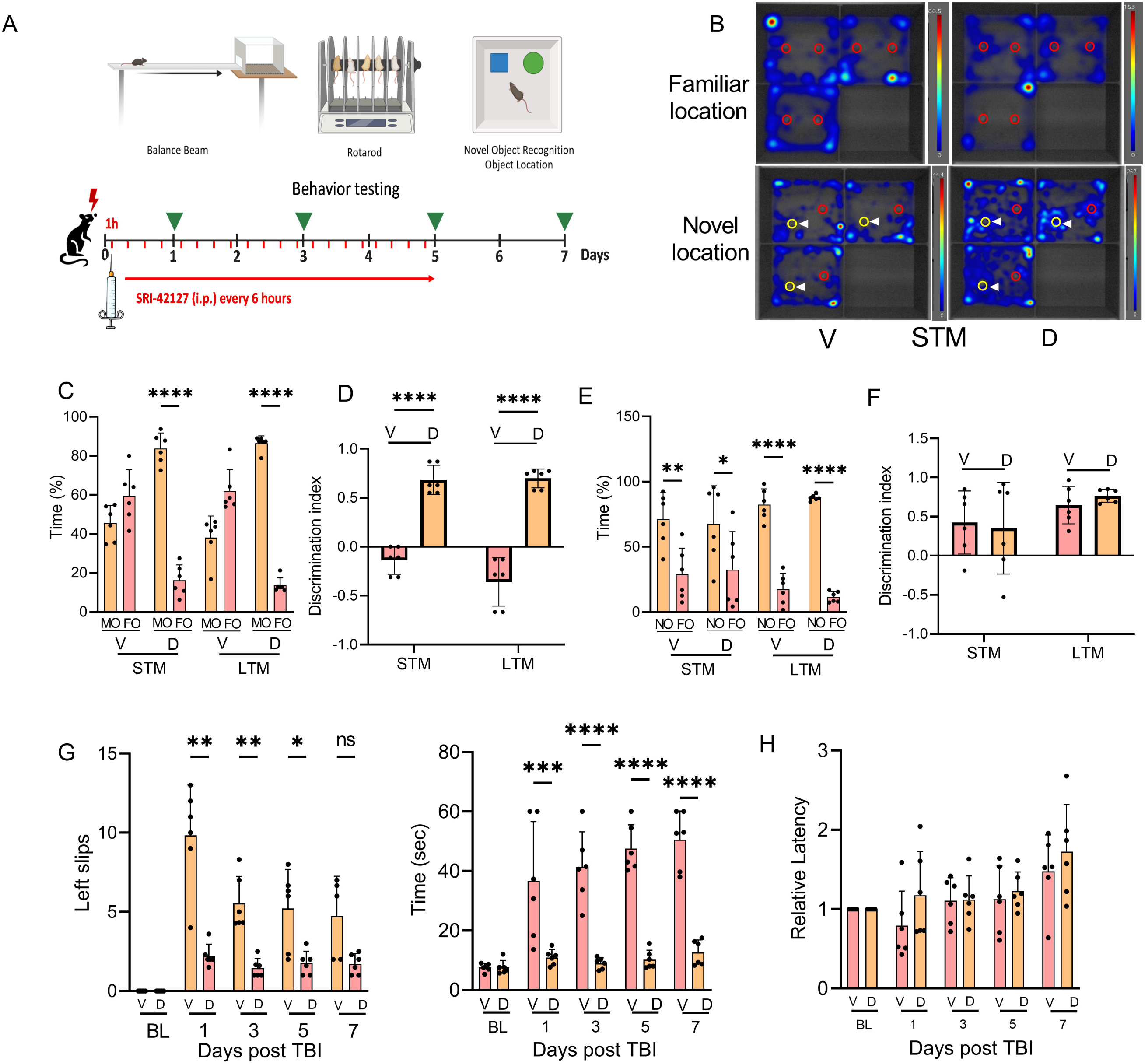
SRI improves memory and motor outcomes after TBI. **(A)** Schematic of experimental protocol for TBI, SRI-42127 administration, and behavioral testing. Mice underwent CCI at day 0 and received SRI-42127 or vehicle every 6 h for 5 days beginning 1 h post-injury; behavioral testing was performed on days 1, 3, 5, and 7. **(B)** Representative heat maps of exploratory behavior during familiarization and testing phase for short term memory for the object location test. SRI-treated mice (D) showed increased exploration concentrated near the moved object, while vehicle-treated mice (V) showed no clear preference between object locations. White arrowheads/yellow circles identify the moved object at the novel location; red circles mark the familiar (unmoved) object. Color scale is normalized to the maximum exploration time within each individual heat map. **(C, D)** Object location test with % exploration time moved vs familiar object and discrimination index for short– and long-term memory. **(E, F)** novel object recognition test showing % time novel object vs familiar object and discrimination index. **(G)** beam walking test assessing crossing time and left foot slips on days, 1, 3, 5, and 7 after TBI. One outlier at day 7 was identified by Grubbs test (α = 0.05) and excluded. BL, baseline. **(H)** Rotarod testing was performed on days, 1, 3, 5, and 7 post-TBI and latency to fall was measured. Data are the mean ± SD (n=6/group); *p<0.05, **p<0.01, ***p<0.001, **** < 0.0001. D, SRI-42127; FO, familiar object; LTM, long-term memory; MO, moved object; NO, novel object; STM, short-term memory; V, vehicle control.

We next assessed the animals with a beam walking test to measure both the time required by mice to cross the beam and the number of left-foot slips **(Fig. 1G)**. At day 1, SRI-42127-treated mice crossed the beam significantly faster, showing a ∼70% reduction in crossing time compared with vehicle-treated controls (p < 0.001). This improvement persisted throughout the observation period up to day 7. Since the TBI was in the right hemisphere, slips occurred in the left forelimb which was then quantitated by an observer blinded to the experimental conditions. SRI-42127-treated mice displayed a ∼75% reduction in left-sided slips at days 1 and day 3 and ∼66% reduction at day 5. At day 3, improvement was observed in vehicle-treated mice and by day 7 there was no significant difference with the drug-treated group (although a trend toward an increase in slips was still present). Sham mice crossed the beam in a similar time at baseline and 1 day after surgery and showed no left-foot slips at either time point (Suppl. Fig. 1).

This suggests that treatment with SRI-42127 during the acute TBI improves fine motor control. We evaluated rotarod performance and observed that neither group exhibited a reduction in mean latency following injury consistent with the location of the TBI which was limited to the dorsal hippocampus and posterior sensorimotor cortex **(Fig. 1H).** Rotarod performance is predominantly governed by the cerebellar and prefrontal circuits (Scholz et al., 2015). Collectively, these findings indicate that the pattern of behavioral outcomes depends on the specific brain regions targeted by the injury and that SRI-42127 treatment improves both spatial memory and fine motor control following TBI.

### 3.2. SRI-42127 blocks nucleocytoplasmic translocation of HuR in microglia and astrocytes

HuR shuttling from the nucleus to the cytoplasm is essential for its function in stabilizing cytokine mRNAs and enhancing their translation (Guha et al., 2023; Guha et al., 2025). The mechanism of action for SRI-42127 is to block HuR nucleocytoplasmic translocation (Filippova et al., 2021; Filippova et al., 2017). To determine whether this occurred in microglia and astrocytes of mice treated with SRI-42127 after TBI, we quantitated HuR nuclear and cytoplasmic localization at 24 h post injury using Fiji software as previously described by our group (**Fig. 2**) (Chellappan et al., 2022; Husain et al., 2025). With vehicle, there was cytoplasmic translocation of HuR in microglia as indicated by a merged yellow signal with HuR and IBA1 immunostaining (**Fig. 2A**) and astrocytes as indicated by merged yellow signal with HuR + GFAP immunostaining in astrocytes (**Fig. 2B**). Translocation of HuR was attenuated with SRI-42127 treatment in both microglia and astrocytes as indicated by absence of merged signal. Quantitative analysis of the nuclear/cytoplasmic HuR ratio across multiple cells showed a 5.4-fold increase in microglia (P<0.0001) and 5.9-fold increase in astrocytes (P<0.0001) after SRI-42127 treatment consistent with HuR nuclear retention **(Fig.2 C,D).** In summary, these findings indicate prominent HuR cytoplasmic translocation in microglia and astrocytes after TBI which is suppressed with SRI-42127 treatment.

**Fig. 2:**
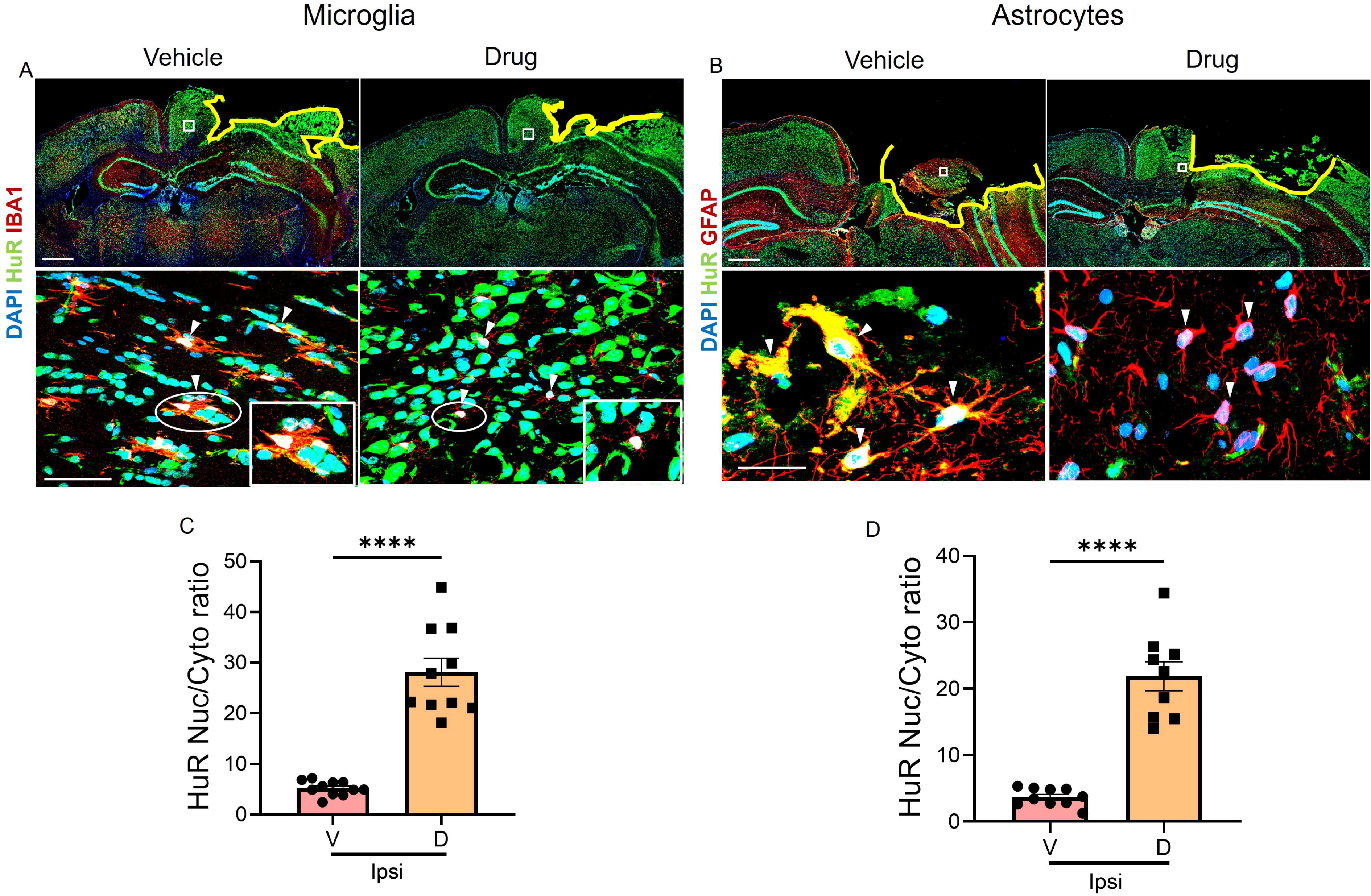
HuR translocates to cytoplasm in microglia and astrocytes in acute TBI. Tissue sections were obtained at 24 h after TBI from vehicle– and SRI-42127-treated mice and double immunostained with antibodies to HuR and Iba1 for microglia **(A)** and HuR/GFAP for astrocytes **(B**). In the upper panels, a yellow line demarcates the lesion core and a white box identifies a ROI in the peri-lesional region for analysis. Scale bars, 1000 μm. The lower panels show HuR translocation in vehicle treated mice as indicated by a merged signal with Iba1 or GFAP (arrows); whereas in SRI-42127 treated mice, no merged signal is observed (arrowheads). Inset is an enlargement of the area highlighted by the ellipse. (**C, D**) HuR localization was quantified for microglia and astrocytes in the peri-lesional area by analyzing 10-11 cells from 4 biological replicates and expressed as a nuclear/cytoplasmic ratio (see Methods). ****p < 0.0001, mean ± SD, scale bar, 10 μm. Cyto, cytoplasm; D, SRI-42127; Nuc, nuclear; V, vehicle control.

### 3.3. SRI-42127 treatment attenuates neuronal loss and reduces the number of apoptotic neurons

Brain sections from ipsilateral hemisphere 24 h after injury were immunostained with NeuN antibodies. The uninjured contralateral hemisphere was assessed for comparison. We observed a ∼ 3.5-fold increase in NeuN+ cells in SRI-42127-treated mice compared to vehicle (P<0.0001) (**Fig. 3A, B**). The NeuN+ cell count approached that of the contralateral hemisphere. No differences in cell counts were observed between SRI-42127 and vehicle-treated mice in the contralateral hemisphere. To evaluate neuronal apoptosis within the perilesional area, sections were co-stained for NeuN and cleaved-Caspase 3 and double-positive cells were quantified (**Fig. 3C, D**). In the vehicle group, a large number of NeuN+/cleaved-Caspase-3+ cells were observed which is consistent with active apoptosis. Conversely, there was a 3-fold reduction in the percentage of apoptotic neurons in SRI-42127-treated mice (P<0.0001). The uninjured hemispheres were negative for immunostaining in both SRI-42127 and the vehicle-treated mice (**Suppl. Fig. 2).** In summary, SRI-42127 treatment led to effective preservation of neurons and a concomitant reduction in apoptosis.

**Fig. 3:**
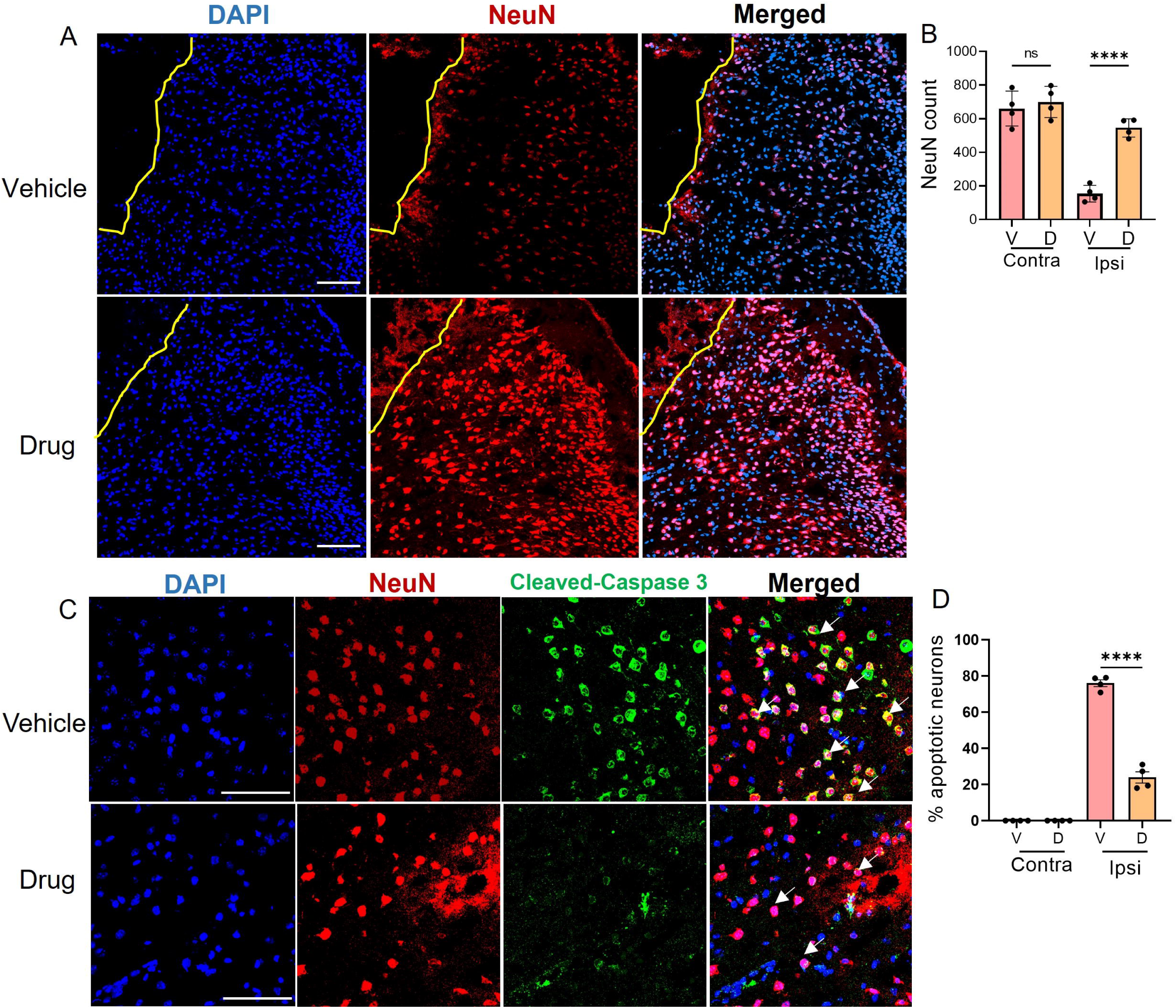
SRI-42127 treatment reduces neuronal loss and neuronal apoptosis post-TBI. (**A**) Brain sections were immunostained for NeuN at 24 h post-TBI. Representative immunostained sections of the peri-lesional cortex show increased NeuN⁺ cells in SRI-42127-treated mice versus vehicle. The yellow line demarcates lesion core. **(B)** Quantification of NeuN⁺ cells in the ipsilateral hemisphere. **(C)** Representative double immunostained sections of peri-lesional cortex with antibodies to NeuN and Cleaved-Caspase 3. The vehicle control exhibits a high number of NeuN+/Cleaved-Caspase 3+ cells (white arrows), reflecting a high number of neurons undergoing apoptosis versus SRI-42127 treated mice. **(D)** Percentage of NeuN⁺ neurons co-labeled with cleaved caspase-3. P value, ****< 0.0001, mean ± SD, scale bars 50 μm, 10 μm. n = 4 mice per group. Contra, contralateral (uninjured) hemisphere; D, SRI-42127; Ipsi, ipsilateral (injured) hemisphere; V, vehicle control.

### 3.4. SRI-42127 reduces neutrophil infiltration in the brain after TBI

Following TBI, a robust inflammatory response occurs with neutrophils being one of the earliest peripheral immune cells to infiltrate the injured brain, peaking within the initial 24 h after injury (Gyoneva and Ransohoff, 2015). Activated neutrophils generate ROS, triggering degranulation and the release of toxic granule enzymes, including myeloperoxidase (MPO) and neutrophil elastase (NE) (Borregaard, 2010). Accordingly, MPO and NE are widely used as quantitative markers of neutrophil infiltration and activation in experimental TBI (Biagas et al., 1992; Semple et al., 2015). We assessed neutrophil infiltration within the peri-lesional area at 24 h post-TBI by double staining immunohistochemistry with MPO and NE antibodies **(Fig. 4)**. A robust increase in double-labeled cells was observed in the injured hemisphere of vehicle treated mice which was blunted by ∼ 3-fold in mice receiving SRI-42127 **(Fig. 4A, B)**. No positive cells were detected in the uninjured hemisphere of either vehicle– and SRI-42127-treated mice **(Fig. 4B and Suppl. Fig. 3)**.

**Fig. 4:**
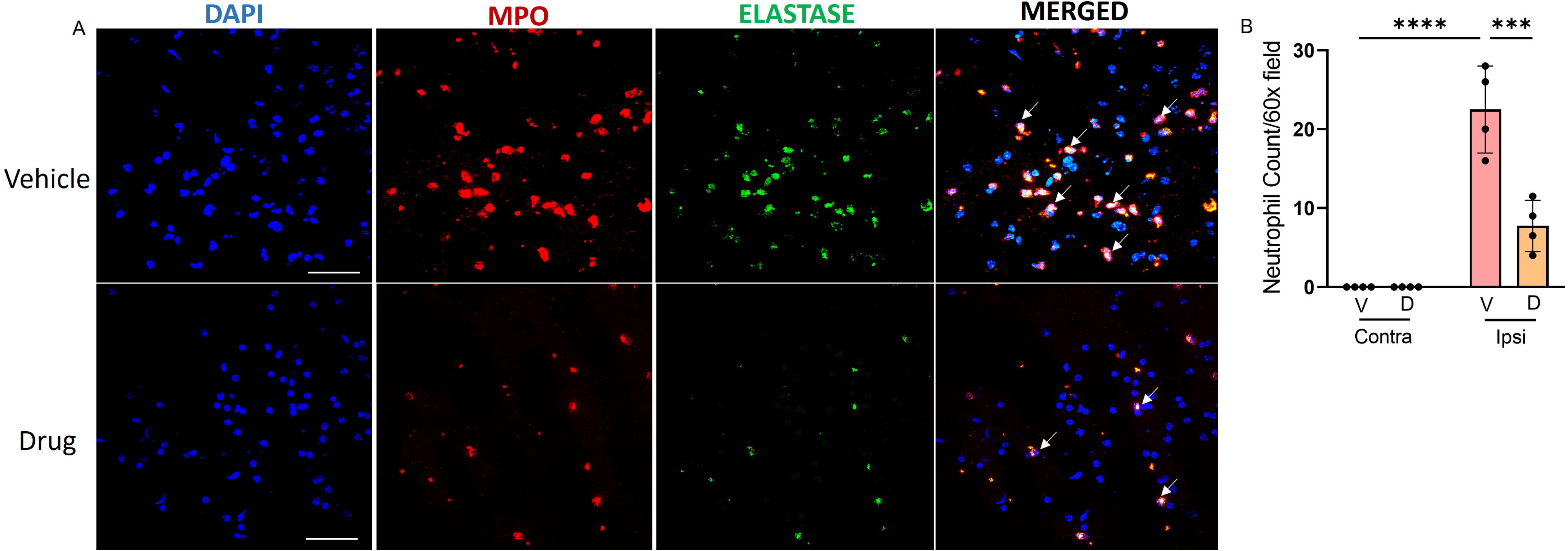
SRI-42127 suppresses neutrophils infiltration 24 h post-TBI. Brain sections of perilesional cortex from SRI-42127– and vehicle-treated mice were double immunostained with MPO and Elastase antibodies. **(A)** Representative images of a drug-treated and vehicle control mouse. MPO+/Elastase+ cells (white arrows) are abundant in the vehicle control mouse. **(B)** MPO+/Elastase+ cells were quantified and show a significant suppression of neutrophil infiltration in SRI-42127-treated mice. P values, ****< 0.0001, ***<0.001, mean ± SD scale bars 10 μm. n = 4 mice per group. Contra, contralateral (uninjured) hemisphere; D, SRI-42127; Ipsi, ipsilateral (injured) hemisphere; V, vehicle control.

### 3.5. SRI-42127 attenuates activation-associated microglia and astrocyte morphological changes

During the early stages of activation, microglia can become hypertrophic, with soma enlargement and increased branch complexity or hyper-ramification (Ziebell et al., 2015). Astrocytes likewise become hypertrophic during reactive astrogliosis, with soma enlargement, complex longer processes, and increased GFAP expression (Sofroniew and Vinters, 2010; Wilhelmsson et al., 2006). These changes are detectable within the first day after injury (Dietrich et al., 1999; Zamanian et al., 2012). We evaluated morphological changes of microglia in the perilesional ipsilateral region at 24 h post-TBI to assess their activation. Using Fiji software, a skeleton analysis of microglia was done as previously described (Green et al., 2022; Husain et al., 2025) (**Fig 5A, B**). In SRI-42127-treated mice, we observed a decrease in branch length (P<0.0001) and number of branches (P<0.0001) consistent with reduced early-stage hyper-ramification (Reddaway et al., 2023). In addition to hyper-ramification, we measured soma size as an indicator of reactive microglial hypertrophy and found a significant reduction in SRI-42127-treated mice (**Fig. 5C, D**) (P<0.0001). Consistent with this finding, Iba1 fluorescence intensity (FI) for individual microglial cells (11-15 cells) was approximately 3-fold reduced (P=0.0002) in SRI-42127-treated mice **(Suppl. Fig. 4A)**.

**Figure 5:**
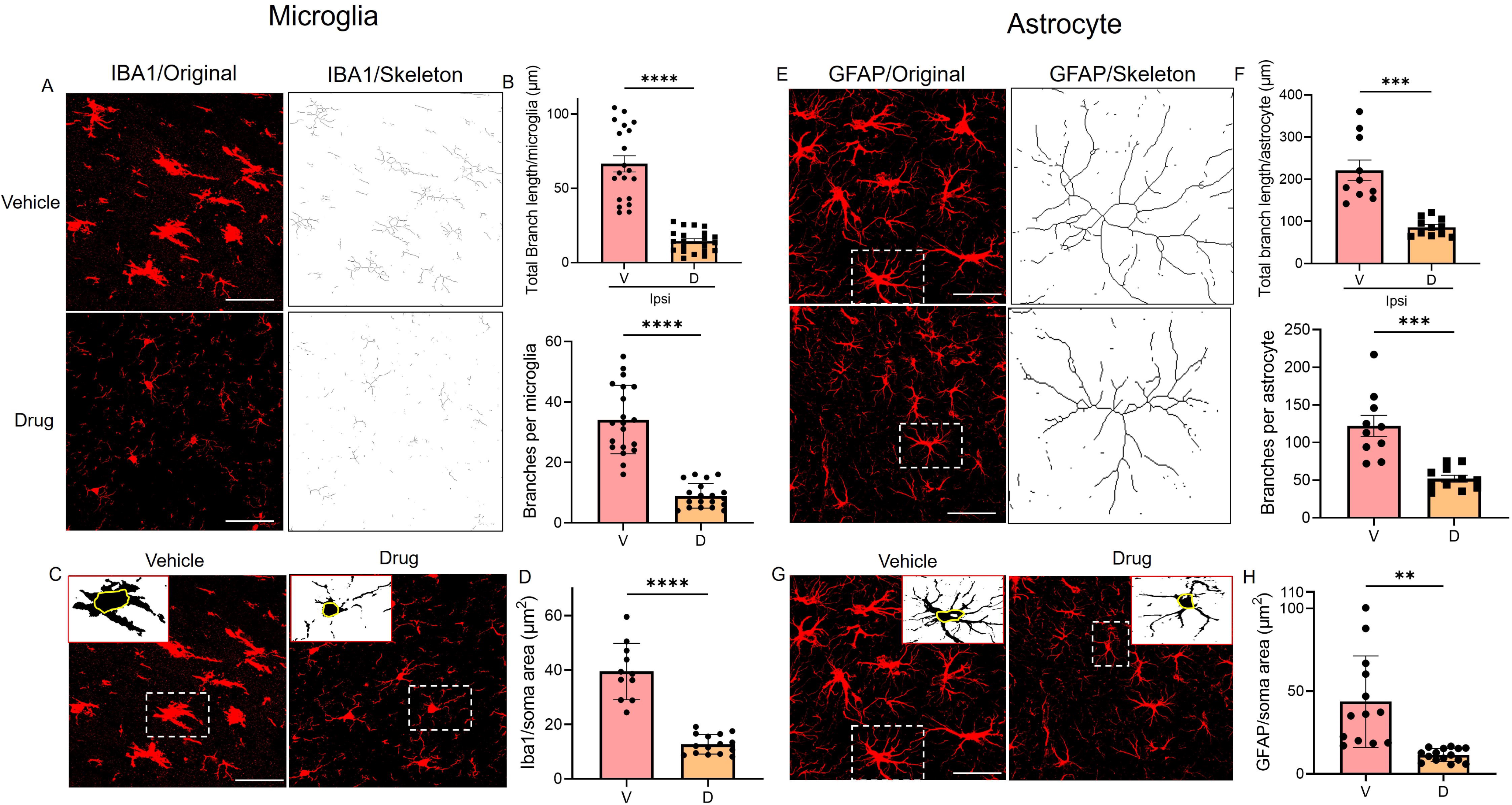
SRI-42127 attenuates activation-associated changes in microglial and astrocyte morphology after TBI. (**A, B**) 24 h after TBI, tissue sections were immunostained with an Iba1 antibody and digitally converted to skeletonized images using Fiji software. Representative images are shown. Morphological features were quantified including total microglia branch length, and number of branches per microglial cell from 20-21 cells (n = 4 biological replicates). **(C)** The soma of Iba1+ microglial cells were identified as shown in the representative images. Binary images were generated by adjusting the Fiji threshold level (shown in inserts). **(D)** Total area was quantified in 11-15 cells from 4 biological replicates. **(E, F)** 24 h after TBI, tissue sections were immunostained with a GFAP antibody and digitally converted to skeletonized images using Fiji software. Representative images are shown. Morphological features were quantified including total astrocyte branch length, and number of branches per astrocyte from 10-11 cells (n = 4 biological replicates). **(G,H)** Astrocyte soma area was quantified in 11-15 cells (n = 4 biological replicates). Error bars represent SD. P values, **** < .0001, ***<0.001, **<0.01. D, SRI-42127; V, vehicle control.

Applying the same skeleton analysis used for microglia, we found that astrocytes in SRI-42127-treated mice had decreased branch length (P=0.0003), number of branches (P=0.0006), and smaller soma size (P=0.0012) (**Fig. 5E-H**). GFAP FI per astrocyte was significantly higher in the perilesional ipsilateral cortex than in the contralateral cortex at 24 h (P<0.0001), indicating injury-induced astrogliosis. SRI-42127 reduced ipsilateral GFAP FI (P<0.0001), whereas neither Iba1 nor GFAP FI differed between vehicle or drug on the contralateral side, confirming that the effect was restricted to the injured hemisphere **(Suppl. Fig. 4B)**. Taken together, SRI-42127 attenuates microglial and astrocyte activation in the early stage of TBI.

### 3.6. SRI-42127 attenuates lesion size, edema and BBB permeability in the acute phase of TBI

T2-weighted MRI lesions in the acute phase of TBI reflect edema and tissue injury (Sanchez-Molano et al., 2023) and can be quantitated volumetrically to measure the extent of edema. We used this modality to assess edema volume at 24 h after SRI-42127 treatment (**Fig. 6A**). With SRI-42127 treatment, we observed a 2.6-fold reduction in lesion volume (21.16 mm³ to 7.93 mm³; P=0.007) and 3-fold reduction in edema volume (from 22.39 mm³ to 7.45 mm³; P=0.006) compared to vehicle (**Fig. 6B**). In support of this finding, brain water content percentage in the injured hemisphere significantly decreased in SRI-42127-treated mice compared to control (79.6 ± 1 versus 83.8 ±0.7; *P* =0.0002) **(Fig. 6C).** We next measured lesion size histologically with cresyl violet staining of brain sections through the injured region (**Fig. 6 D, E**). Lesion size was significantly reduced in SRI-42127-treated mice by ∼2-fold compared to vehicle (P <0.0001). We extended the imaging analysis to 7 d post injury and observed a similar pattern of reduced lesion volume (P =0.032) and a trend toward reduced edema (P =0.095) in SRI-42127-treated mice (**Suppl. Fig. 5A, B**). We next assessed BBB integrity at 24 h post injury with Evans blue dye (**Fig. 6 F, G**). To quantitate dye extravasation into brain tissue, spectrophotometric OD values were measured in the injured hemisphere (μg/g) and normalized to the uninjured hemisphere to account for variability between groups. SRI-42127 treatment resulted in a ∼3.9-fold reduction in dye content (P < 0.0001) compared to vehicle. In the uninjured hemispheres, there was no difference between drug– and vehicle-treated groups. Taken together, SRI-42127 blunted post-injury edema and BBB disruption resulting in a significantly reduced lesion size.

**Fig. 6:**
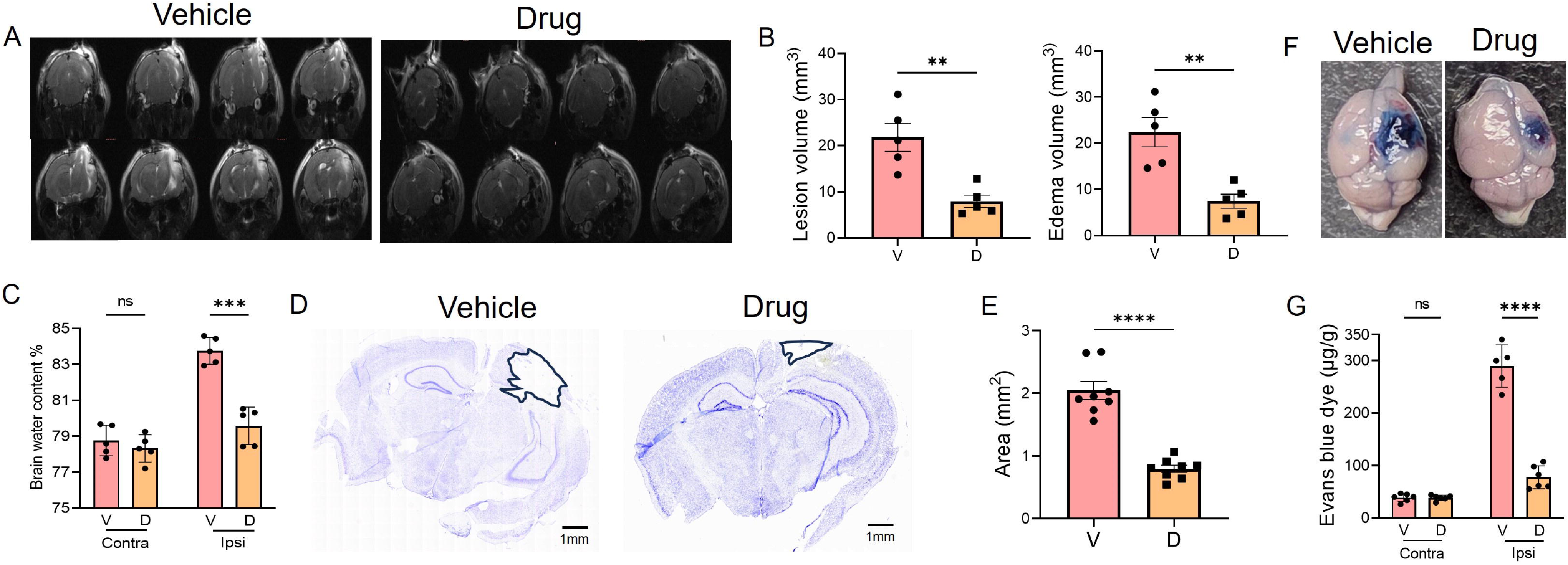
SRI-42127 suppresses edema, lesion size and reduces BBB damage 24 h after TBI. (**A**) Representative coronal T2-weighted MRI images of a vehicle– or SRI-42127-treated mouse at 24 h post-TBI. (**B**) MRI Lesion volume and edema were quantitated for 5 vehicle– and 5 SRI-42127-treated mice. **(C)** Brain water content (%) in the contralateral and ipsilateral hemispheres of vehicle– and SRI-42127-treated mice. SRI-42127 significantly reduced ipsilateral brain water content compared to vehicle, with no difference between groups in the contralateral hemisphere. Data points represent the mean ± SD of 5 biological replicates. P value: ***< 0.001. **(D)** Representative cresyl violet-stained brain sections from a vehicle-treated or SRI-42127-treated mouse at 24 h post-TBI. Injured areas lack staining and are outlined in black. (**E**) Lesion area was quantified (n = 4 biological replicates). **(F)** Representative whole brain image of Evans blue dye staining for a vehicle– and SRI-42127-treated mouse. **(G)** Evans blue dye content was quantified in brain tissue from the injured and uninjured hemispheres in 5-6 biological replicates. Error bars represent SD. P values, **** < .0001, **<0.01. Contra, contralateral (uninjured) hemisphere; D, SRI-42127; Ipsi, ipsilateral (injured) hemisphere; V, vehicle control.

### 3.7. HuR inhibition by SRI-42127 blunts the induction of proinflammatory mediators after CCI

At 24 h post TBI, tissue samples were obtained from the lesion site and corresponding contralateral (uninjured) side to assess induction of proinflammatory mediators. We observed a 2-to 50-fold induction of major proinflammatory mediator mRNAs, including *IL-1*β, *IL-6*, *TNF-*α, *CXCL1*, *CXCL2*, *CCL2*, *MMP12*, *iNOS*, *CCR2*, *NLRP3*, and *NOX2,* on the injured side compared to uninjured side in vehicle-treated mice (**Fig. 7A**). The induction was most pronounced with *CXCL2* (∼50-fold) and *MMP12* (∼39-fold). SRI-42127 treatment suppressed each of these proinflammatory mediators, most potently for *CXCL2* (∼21-fold) and *MMP12* (∼18-fold). Other mRNAs were suppressed by ∼1.5-to 9-fold. *iNOS* trended toward suppression but did not reach significance due to higher variability of mRNA levels. *COX2* mRNA was not induced or affected by SRI-42127 treatment. Given our previous observation that HuR affects COX-2 protein expression rather than the mRNA after neurotrauma (Husain et al., 2025), we assessed expression by western blot **(Fig. 7B and Suppl. Fig. 6).** We observed a ∼2.4-fold increase in COX-2 protein levels on the injured side compared to the contralateral side in vehicle-treated mice. With SRI-42127 treatment, there was a 2.8-fold (P=0.007) reduction in COX-2 expression to a level similar to the uninjured side. To determine the impact of SRI-42127 on inflammatory cytokine induction at a later timepoint, we assessed brains at day 7 post injury which is 2 days after SRI-42127 treatment ended (**Suppl. Fig. 7**). Induction of proinflammatory mediators persisted in the vehicle-treated mice and ranged from 2-to 60-fold versus the uninjured side although there was variability within the groups. The fold-induction of *MMP-12*, *TNF-*α, *NOX2*, *NLRP3*, and *CCR2* was even higher than at day 1 whereas *CXCL1* and *CXCL2* were lower. *IL-1*β, *CCL2,* and *iNOS* showed similar changes. Interestingly, *IL-6* induction was no longer detected. To validate use of the contralateral hemisphere as an internal control, sham mice were compared to vehicle– and drug-treated injured mice for *IL-6* induction at 1 day. We observed similar expression levels of *IL-6* in sham control compared to the contralateral hemisphere of injured vehicle– and drug-treated mice (**Suppl. Fig. 8**). There was a ∼20-fold induction of *IL-6* in the ipsilateral injured hemisphere of vehicle-treated mice which was suppressed nearly back to baseline with SRI-42127 treatment. Taken together, treatment with SRI-42127 in the acute phase of TBI broadly suppressed the induction of proinflammatory mediators at the level of injury and this persisted in the subacute phase after SRI-42127 treatment ended.

**Figure 7:**
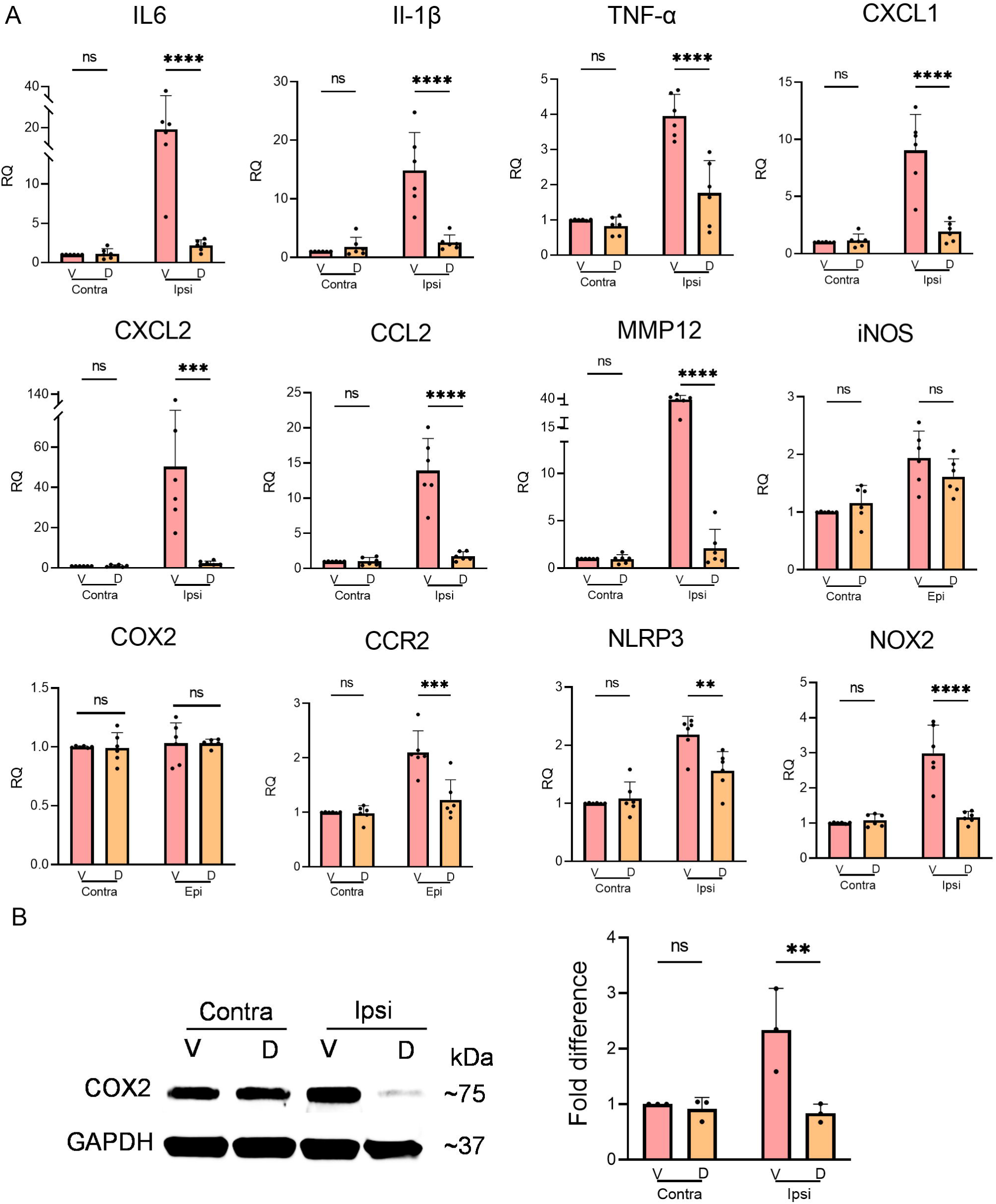
SRI-42127 attenuates proinflammatory cytokine and chemokine induction after TBI. (**A)** RNA was extracted from brain tissue at 24 h post-TBI in the ipsilateral and contralateral hemispheres. RNA expression levels of proinflammatory mediators were assessed by RT-qPCR. All values were expressed relative to the contralateral hemisphere of the vehicle control which was set at 1. *HPRT* was used as the housekeeping control. P values: * < 0.05, ** < 0.01, *** < 0.001, **** < 0.0001. Error bars represent means ± SD of 6 mice/group. **(B)** COX-2 immunoblot of protein lysate from Ipsilateral and contralateral brain tissue 24 h after TBI. A representative blot is shown. COX-2 levels were quantified by densitometry from three biological replicates. All values are expressed relative to the contralateral side of the vehicle control which was set at 1. Error bars represent the mean ± SD. P values: **< 0.01. Contra, contralateral (uninjured) hemisphere; D, SRI-42127; Ipsi, ipsilateral (injured) hemisphere; V, vehicle control.

## 4. Discussion

The primary injury of TBI triggers a cascade of proinflammatory responses, initiated by microglia and astrocytes that are rapidly activated by the mechanical strain and exposure to damage-associated molecular patterns (DAMPs), and then amplified by infiltrating peripheral immune cells (Burda et al., 2016; Davis and Godbout, 2026; Simon et al., 2017). This cascade results in the induction and release of diverse proinflammatory mediators which promote secondary injury from BBB disruption, edema, ischemia, and direct cytotoxicity (Jha et al., 2024; Zhang et al., 2025; Zhu et al., 2026). The diversity of these inflammatory mediators (produced by many cell types) and their overlap in driving pathways of secondary tissue injury provide a plausible explanation as to why therapies targeting neuroinflammation have not been successful (Jha et al., 2024). In this report, we show that inhibiting HuR, an RNA regulator of proinflammatory mediators expressed in glial cells and most peripheral immune cells, can overcome these limitations and potently suppress post-TBI neuroinflammation and secondary tissue injury (Guha et al., 2025; Guha et al., 2022; Majunder et al., 2022; Wang et al., 2019).

In our previous work, we found that modulating HuR in microglia and astrocytes, either by silencing or upregulating, reduced or increased (respectively) the expression of many proinflammatory mediators that drive secondary injury after TBI including IL1β, IL-6, and TNF-α, (Chellappan et al., 2022; Davis and Godbout, 2026; Kwan et al., 2017a; Kwan et al., 2017b; Matsye et al., 2017; Wang et al., 2019). This effect has been observed more broadly in other immune cells in peripheral inflammatory disorders and cancer, providing a rationale for investigating the impact of inhibiting HuR in TBI (Guha et al., 2022; Majunder et al., 2022; Schultz et al., 2020; Wang et al., 2019). A key signature of HuR activation is nucleocytoplasmic translocation, a process essential for its ability to stabilize and enhance the translational efficiency of bound proinflammatory mRNAs via AREs in the 3’ UTR (Fan and Steitz, 1998; Guha et al., 2023; Meisner et al., 2007; Nabors et al., 2001). In the current study, HuR translocation was evident in microglia and astrocytes of vehicle-treated mice at 24 hours post TBI consistent with its activation. We observed a similar pattern in astrocytes and microglia after SCI suggesting that glial HuR translocation is a more generalized response to acute CNS injury (Husain et al., 2025; Kwan et al., 2017a; Kwan et al., 2017b). The inhibitor used in this study, SRI-42127, was identified by our group in a high throughput screen for molecules that block HuR homodimerization (Filippova et al., 2021; Filippova et al., 2017), a process required for its nucleocytoplasmic translocation (Meisner et al., 2007). Inhibition of HuR translocation after SRI-42127 treatment was evident in astrocytes and microglia 24 hours post TBI similar to what we observed in SCI (Husain et al., 2025) and in a LPS model of CNS neuroinflammation (Chellappan et al., 2022), reflecting the consistency of this drug effect.

The impact of SRI-42127-induced HuR inhibition after TBI was a marked attenuation of the proinflammatory milieu in the injured brain as reflected by the broad suppression of proinflammatory mediators. This included major drivers of secondary injury such as IL-1β, IL-6, and TNF-α (Wofford et al., 2019; Zhu et al., 2026). Microglial/astrocyte activation, BBB disruption, and neutrophil infiltration are three key pathways that contribute to this proinflammatory microenvironment (Burda et al., 2016; Jassam et al., 2017). Although the relative contribution of each pathway cannot be discerned due to their convergence in this early phase, all were attenuated by SRI-42127. For the first pathway, microglia and astrocytes become activated within minutes to hours after injury, as gauged by morphological changes, and are considered initiators of the proinflammatory cascade (Burda et al., 2016; Davalos et al., 2005; Jassam et al., 2017). Based on our prior work, HuR plays a major role in the broad induction of proinflammatory mediators in activated microglia and astrocytes and this can be rapidly reversed with SRI-42127 or HuR silencing (Chellappan et al., 2022; Husain et al., 2025; Kwan et al., 2017a; Matsye et al., 2017). A key factor in the potency of SRI-42127 for quelling early neuroinflammatory responses is related to its pharmacological properties. After peripheral administration, SRI-42127 quickly penetrates the CNS and reaches therapeutic levels within minutes after peripheral administration (even with an intact BBB) (Filippova et al., 2021). The rapid action of SRI-42127 was evident in a prior study of LPS-induced CNS neuroinflammation where there was robust repression of proinflammatory mediators, including IL-1β, IL-6, TNF-α and CXCL1, in all brain regions with a single dose of SRI-42127 two hours after LPS injection (Chellappan et al., 2022). These properties would enable SRI-42127 to inhibit early triggers of the inflammatory cascade after TBI and the “snow ball” amplification effect that ensues. The ease of administration is favorable for early clinical intervention, potentially even by emergency personnel.

The second pathway that was blunted by SRI-42127 was disruption of the BBB as reflected by suppression of Evans blue dye extravasation, edema by MRI testing, and lower brain water content. The proinflammatory environment including IL-6, TNF-α, IL-1β, and CCL2, ties directly into early BBB disruption and increased permeability observed after TBI (Cash and Theus, 2020; Semple et al., 2010). The induction of *CCL2* and its signaling receptor, *CCR2*, was essentially abrogated as we observed previously with SCI (Husain et al., 2025). Astrocytes are major producers of CCL2 which is released by end-feet processes abutting vascular endothelium, leading to increased permeability via CCR2-mediated signaling in vascular endothelial cells (Chodobski et al., 2011; Semple et al., 2010). This signaling axis may be especially relevant as *CCR2* is also regulated by HuR (Matsye et al., 2017) and was markedly suppressed by SRI-42127 in this study and our prior study with SCI (Husain et al., 2025). Of note, HuR is expressed in vascular endothelial cells and regulates induction of inflammatory responses and thus may have been suppressed by SRI-42127 treatment (Rhee et al., 2010; Si et al., 2021).

MMPs also play a critical role in BBB disruption by degrading the vascular basement membrane and extracellular matrix which support the integrity of tight junctions (Jassam et al., 2017). *MMP12* was induced by ∼40-fold at 24 hours and 60-fold at 7 days after TBI and suppressed nearly back to baseline with SRI-42127. This is consistent with our prior report that HuR strongly regulates *MMP12* expression in activated microglia and astrocytes (Kwan et al., 2017a; Matsye et al., 2017). MMP12 has been shown to play a direct role in BBB disruption after ischemic stroke by degrading tight junction proteins (Chelluboina et al., 2015). It also regulates MMP-9 which is a recognized disruptor of BBB integrity after TBI (Sunny et al., 2024). Of note, *MMP12* was persistently induced to high levels at 7 days post injury suggesting that it may contribute to BBB dysfunction in subacute phases of TBI or longer. The striking suppression of *MMP12* by SRI-42127 also persisted. Further studies will be required to assess the longevity of this suppressive effect and the long term impact on BBB integrity.

Another critical driver of BBB disruption and edema is oxidative stress (Fesharaki-Zadeh, 2022). Two key enzymes in the generation of deleterious reactive oxygen species (ROS) after TBI are NOX2 and COX-2, and either the mRNA (*NOX2*) or protein (COX-2) induction was essentially blocked by SRI-42127 (Bano et al., 2025; Dohi et al., 2010; Fang et al., 2022; Hiskens et al., 2024; Kumar et al., 2016). Microglia, astrocytes, and infiltrating macrophages and neutrophils appear to be major producers of these enzymes after TBI and they peak in the initial 24-48 hours. The mitigating effect of suppressing NOX2 in TBI was previously shown where genetic deletion or pharmacological inhibition attenuated microglial/macrophage activation, reduced apoptosis and lesion volume, and improved functional outcomes (Dohi et al., 2010; Kumar et al., 2016). Vascular endothelial cells are also key contributors to ROS generation after TBI (Lutton et al., 2019) and a prior study found HuR to regulate NOX2 in those cells (Si et al., 2021). Interestingly, for COX-2, the protein but not mRNA was induced after TBI reflecting a dissociation between RNA and protein expression that was observed in our prior study in SCI (Husain et al., 2025). This highlights the dual and distinct roles that HuR plays in post-transcriptional regulation: RNA stability and translational efficiency (Guha et al., 2023).

The third pathway that was blunted by SRI-42127 was the infiltration of neutrophils which was suppressed by nearly 3-fold. Neutrophils are the earliest peripheral immune cells to invade the injured brain after TBI, occurring within hours and typically peaking at 24 hours (Gyoneva and Ransohoff, 2015). Several findings from this report provide plausible mechanisms for this suppression. First, BBB disruption facilitates neutrophil transmigration after TBI (Chodobski et al., 2003; Soares et al., 1995) and this was markedly attenuated by SRI-42127. Second, induction of the two major chemokines for neutrophil recruitment, *CXCL1* and *CXCL2*, was strikingly attenuated with expression levels nearing those of the contralateral (uninjured) side (Liu et al., 2018; Petri and Sanz, 2018; Soares et al., 1995). A similar degree of suppression for these chemokines was observed after SCI in mice treated with SRI-42127 (Husain et al., 2025). Astrocytes are a predominant cell type expressing CXCL1 in TBI (Jiang et al., 2023; Xia et al., 2022). Astrocyte activation in TBI is initially triggered by mechanical forces from the primary injury which can initiate a proinflammatory phenotype (Burda et al., 2016). In our previous work, we observed astrocyte activation after mechanical stretch using an *in cellulo* model that mimics acceleration/deceleration forces found in human TBI (Floyd et al., 2001; Kwan et al., 2017a). HuR translocation was observed in stretched astrocytes along with a significant induction of *CXCL1*, *IL-6*, *TNF-*α, *IL-1*β, and *CCL2*. This induction was blocked with HuR silencing or inhibition (Kwan et al., 2017a; Meisner et al., 2007). The importance of astrocytic HuR in driving neuroinflammatory responses relevant to TBI was previously demonstrated in models of stroke (Ardelt et al., 2017) and SCI (Kwan et al., 2017b) where its selective transgenic expression led to disruption of BBB, blood-spinal cord barrier, and edema in the acute phase. It should be noted that other cell types produce CXCL1 and CXCL2 after TBI, such as microglia oligodendrocytes, choroid plexus epithelium, endothelial cells, neurons and infiltrating immune cells (Fang et al., 2022; Jassam et al., 2017; Jiang et al., 2023; Strauss et al., 2000; Szmydynger-Chodobska et al., 2009; Wu et al., 2026; Xia et al., 2022; Zhao et al., 2023). As HuR is expressed in these cell types, the profound suppression observed with SRI-42127 may reflect a collective inhibition of these cells. This possibility was underscored in our study of LPS-induced CNS neuroinflammation where SRI-42127 blocked neutrophil and monocyte infiltration into all brain regions but microglial-specific HuR deletion led to suppression of monocytes only (Chellappan et al., 2022).

The marked and early attenuation of neutrophils in SRI-42127-treated mice at 24 hours post TBI likely played a significant role in reducing secondary injury and improving functional outcome. There is strong evidence that neutrophils are pathogenic in the acute phase and that blocking their recruitment or depleting them after TBI mitigates injury (Liu et al., 2026; Zhao et al., 2023). Evidence from human and preclinical studies show that neutrophils exert cytotoxicity after TBI by forming neutrophil extracellular traps (NETs) (Liu et al., 2026; Shi et al., 2023; Vaibhav et al., 2020). Inhibiting NETs or promoting their degradation reduces microglial activation, BBB injury, edema, and neuronal apoptosis (Liu et al., 2026; Mi et al., 2023; Shi et al., 2023; Slough et al., 2025; Vaibhav et al., 2020). NLRP3 plays an important role in the genesis of NETs and is robustly induced after TBI (Leal and Weber, 2026; Ma et al., 2017; Munzer et al., 2021). Silencing or pharmacologically inhibiting it significantly reduces post-TBI NET formation and secondary injury (Ismael et al., 2020; Liu et al., 2026). SRI-42127 suppressed the induction of *NLRP3* which is consistent with a prior report showing that HuR positively regulates NLRP3 expression through ARE-mediated mechanisms (Wang et al., 2024). Another contributing factor may have been the concomitant suppression of NOX2 which is an upstream regulator of NLRP3 required for its induction in the acute phase of TBI (Laabei et al., 2025; Ma et al., 2017).

The net effect of HuR inhibition with SRI-42127 was reduced lesion size and increased neuronal survival which translated into improved cognitive and motor function in the short term. Deficits in hippocampal-dependent spatial memory as reflected by the object location test is consistent with the injury being centered on the dorsal hippocampus and posterior sensorimotor cortex (Paterno et al., 2018). The anatomical specificity was underscored by the preservation of object recognition memory in both groups, which relies more heavily on perirhinal cortex than on an intact hippocampus (Broadbent et al., 2010). Fine motor control was also improved, with faster beam crossing and fewer foot slips in the days immediately after injury, whereas rotarod performance, which depends more heavily on cerebellar and prefrontal circuits outside the injury area, was unaffected by either injury or treatment.

The ongoing induction of proinflammatory mediators at 7 days post injury in vehicle-treated mice is consistent with progression into more chronic phases of neuroinflammation as previously in the CCI model (Abikenari et al., 2025). Notably, the fold-induction of *MMP12*, TNF-α, *NLRP3*, and *NOX2* was even higher than at 24 hours but continued to be suppressed to control levels with SRI-42127 treatment. It is likely that this persistent suppression of proinflammatory mediators reflects a combination of ongoing HuR inhibition and the early blockade of the inflammatory cascade. Since the NOX2-ROS-NLRP3 inflammasome axis plays a key role in promoting a proinflammatory phenotype in microglia/macrophages in the chronic phase of TBI, the potent effect of SRI-42127 may reduce long-term progression of neurodegeneration and cognitive dysfunction (Henry et al., 2020; Tan et al., 2021). Future studies will be necessary to determine whether the mitigating effects of SRI-42127 go beyond the subacute phase and whether intervention with SRI-42127 in chronic phases can mitigate subsequent tissue injury, BBB injury, and neurodegeneration. A caveat to ongoing treatment with SRI-42127, however, is the transition that occurs with microglia and macrophages (and astrocytes) to a reparative phenotype, supporting neural plasticity and angiogenesis. Some of the HuR-regulated proinflammatory mediators that are deleterious in the acute phase, such as IL-6, TNF-α, and VEGF, change their roles to support this healing process (Li et al., 2021; Scherbel et al., 1999)

## Acknowledgements

We wish to thank the UAB Animal facility, UAB Shelby core, UAB’s Preclinical Imaging Shared Facility Grant P30CA013148, and the MRI S10 instrumentation grant S10 OD028498-01.

## Authors’ contributions

MAH, and PHK contributed to the conception and design of the study. MAH, BS, RS and OT, performed experiments and acquired data. MAH, PHK, BS, OT, YS, SK, LBN, NF contributed to data analysis and interpretation. MAH and PHK prepared figures and drafted the manuscript. APB, and MB performed MRI experiments and image processing. All authors read and approved the final manuscript.

## Availability of data and materials

Data that support the findings of this study are available on request from the corresponding author. The data are not publicly available due to privacy and/or ethical restrictions.

## Figure Legends

**Supplementary Figure 1:** Craniotomy alone does not impair spatial memory or beam walking in sham mice. (A, B) Short-term object location test in sham mice (V; vehicle-treated), showing % exploration time of the moved vs familiar object (A) and the discrimination index (B). (C) Beam crossing time at baseline (BL) and 1 day after surgery (D) no left-foot slips were observed at either time point. Data are the mean ± SD (n = 4). P values: ** < 0.01. FO, familiar object; MO, moved object.

**Supplementary Figure 2:** Analysis of NeuN and Cleaved-Caspase 3 immunostaining in the uninjured (contralateral) hemisphere. (**A)** Representative images of NeuN immunostaining. **(B)** Representative images of cleaved-Caspase 3 immunostaining showing no evidence of neuronal apoptosis. scale bars 50 μm,10 μm. Sections from 3 other biological replicates in drug– and vehicle-treated groups were also negative for NeuN+/cleaved-caspase 3+ double staining (not shown).

**Supplementary Figure 3:** Analysis of MPO and Elastase expression in the uninjured contralateral hemisphere. Representative immunofluorescence images of the contralateral hemisphere from a vehicle– and SRI-42127-treated mouse immunostained for MPO (red), and Elastase (green) antibodies and counterstained with DAPI. The lack of immunopositive cells confirms the absence of infiltrating neutrophils in the uninjured hemispheres of both groups.

**Supplementary Figure 4:** Microglial and astrocyte activation is reduced in SRI-42127-treated mice. **(A)** Representative Iba1 immunostained images of the perilesional injured cortex from vehicle– and SRI-42127-treated mice. Fluorescence intensity (FI) per cell was quantified in the uninjured contralateral and injured ipsilateral hemispheres (11-14 cells from n = 4 mice per group). **(B)** Representative GFAP-stained images from a vehicle– and SRI-42127-treated mouse. GFAP FI per cell was quantified (12-18 cells from n = 4 mice per group). Means ± SD are shown. P values: ****<0.0001; ns, not significant. Contra, contralateral (uninjured) hemisphere; D, SRI-42127; Ipsi, ipsilateral (injured) hemisphere; V, vehicle control.

**Supplementary Figure 5:** SRI-42127 attenuates lesion size after 7 days TBI. **(A)** Representative coronal T2-weighted images of a vehicle or SRI-treated mouse at 7-day post TBI. (**B**) Lesion volume and edema was quantitated for 3 vehicle and 5 SRI-treated mice. Data points represent the mean ± SD. P value: *< 0.05. Contra, contralateral (uninjured) hemisphere; D, SRI-42127; Ipsi, ipsilateral (injured) hemisphere; V, vehicle control

**Supplementary Figure 6:** Uncropped immunoblot of COX-2 and GAPDH from Fig. 7. Contra, contralateral (uninjured) hemisphere; D, SRI-42127; Ipsi, ipsilateral (injured) hemisphere; V, vehicle control.

**Supplementary Figure 7:** Suppression of proinflammatory cytokines persists at 7 days. At 7 days post-TBI, RNA was extracted from both the ipsilateral and contralateral brain regions of injured mice and mRNA levels of proinflammatory mediators were quantified by RT-qPCR. Outliers were identified by Grubbs test (α = 0.05) and excluded. P values: * < 0.05, **< 0.01, ***< 0.001, ****P<0.0001. Error bars denote the mean ± SD (n=3-6 biological replicates/group).

**Supplementary Figure 8:** Sham controls showed similar *IL-6* levels compared to contralateral hemisphere in injured mice. Relative *IL-6* mRNA expression in brain tissue was assessed by RT-qPCR in the ipsilateral hemisphere of vehicle-treated sham controls (set at 1) and compared to ipsilateral (injured) and contralateral (uninjured) hemispheres of vehicle– and SRI-42127-treated mice. *IL-6* levels were similar between the sham group compared to contralateral hemispheres of vehicle– and drug-treated mice. There was significant *IL-6* induction in the ipsilateral hemisphere of vehicle-treated injured mice which was suppressed by SRI-42127. Data are the mean ± SD (n=5-6/group); ****P<0.0001; ns, not significant. Contra, contralateral (uninjured) hemisphere; D, SRI-42127; Ipsi, ipsilateral (injured) hemisphere; RQ, relative quantity; V, vehicle control.

