## Supplementary figures and images for "Targeting the RNA regulator HuR in acute traumatic brain injury mitigates secondary injury and improves functional outcome by suppressing neuroinflammation"

### Suppl fig 1

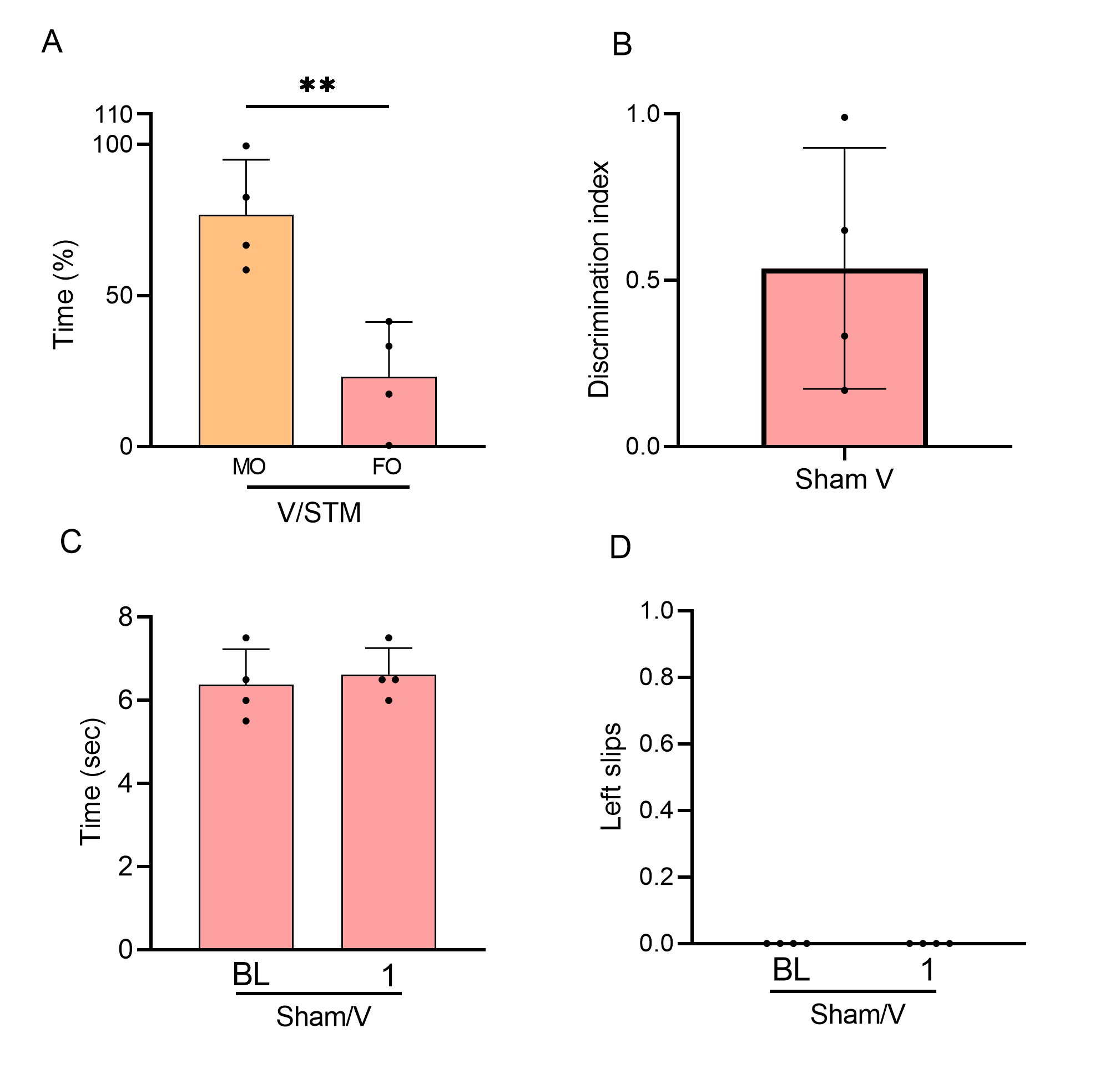

### Suppl fig 2

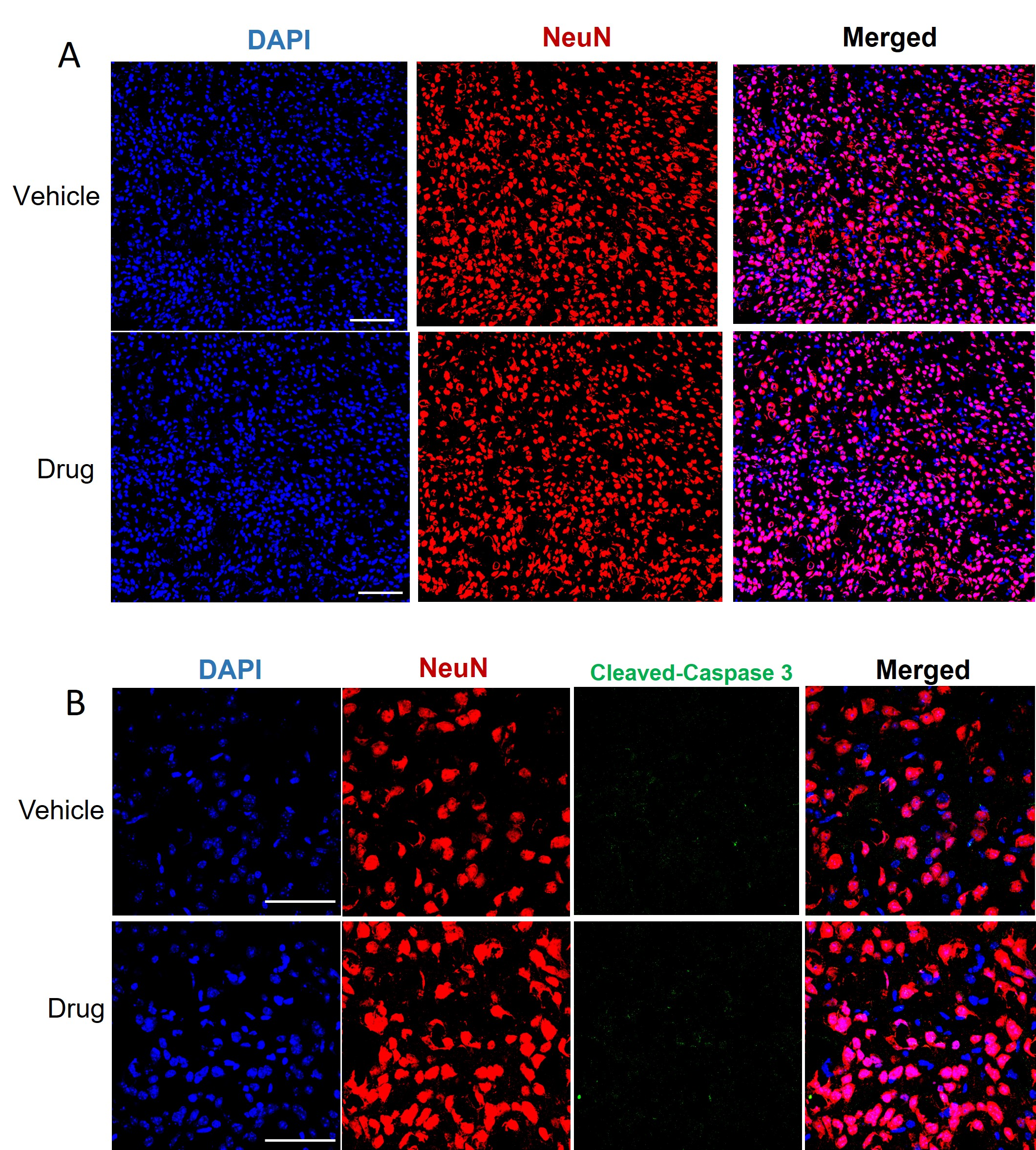

### Suppl fig 3

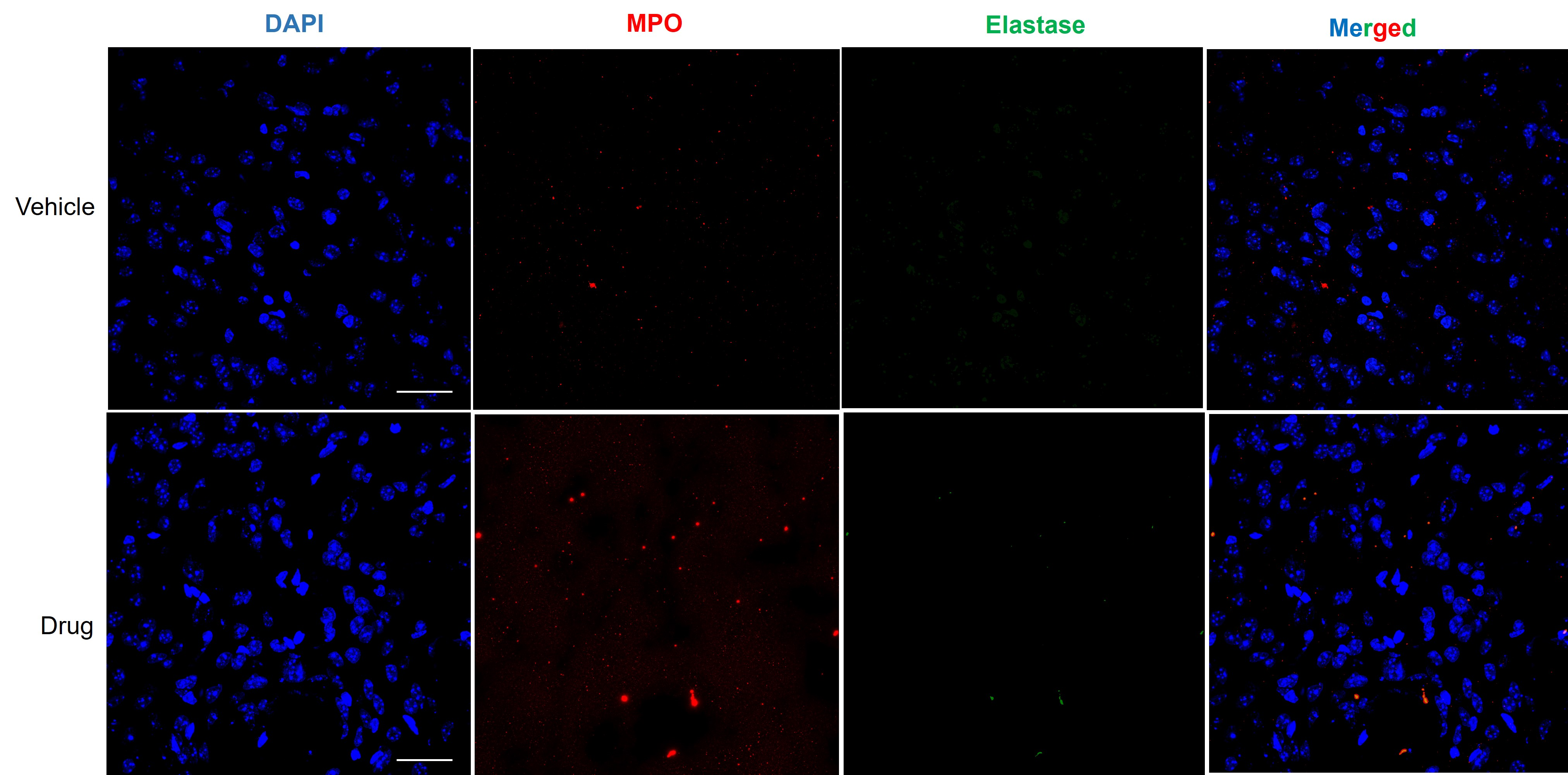

### Suppl fig 4

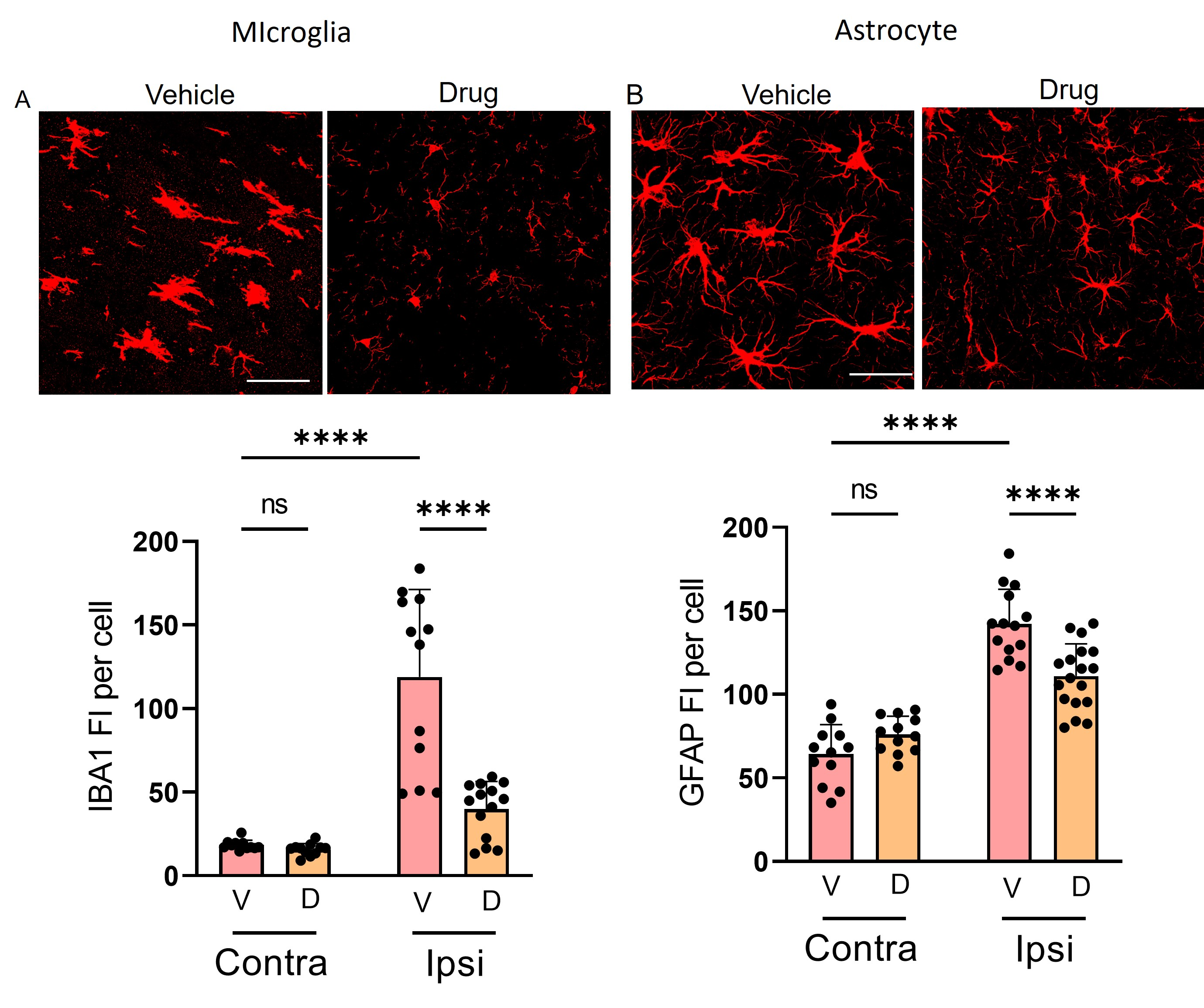

### Suppl fig 5

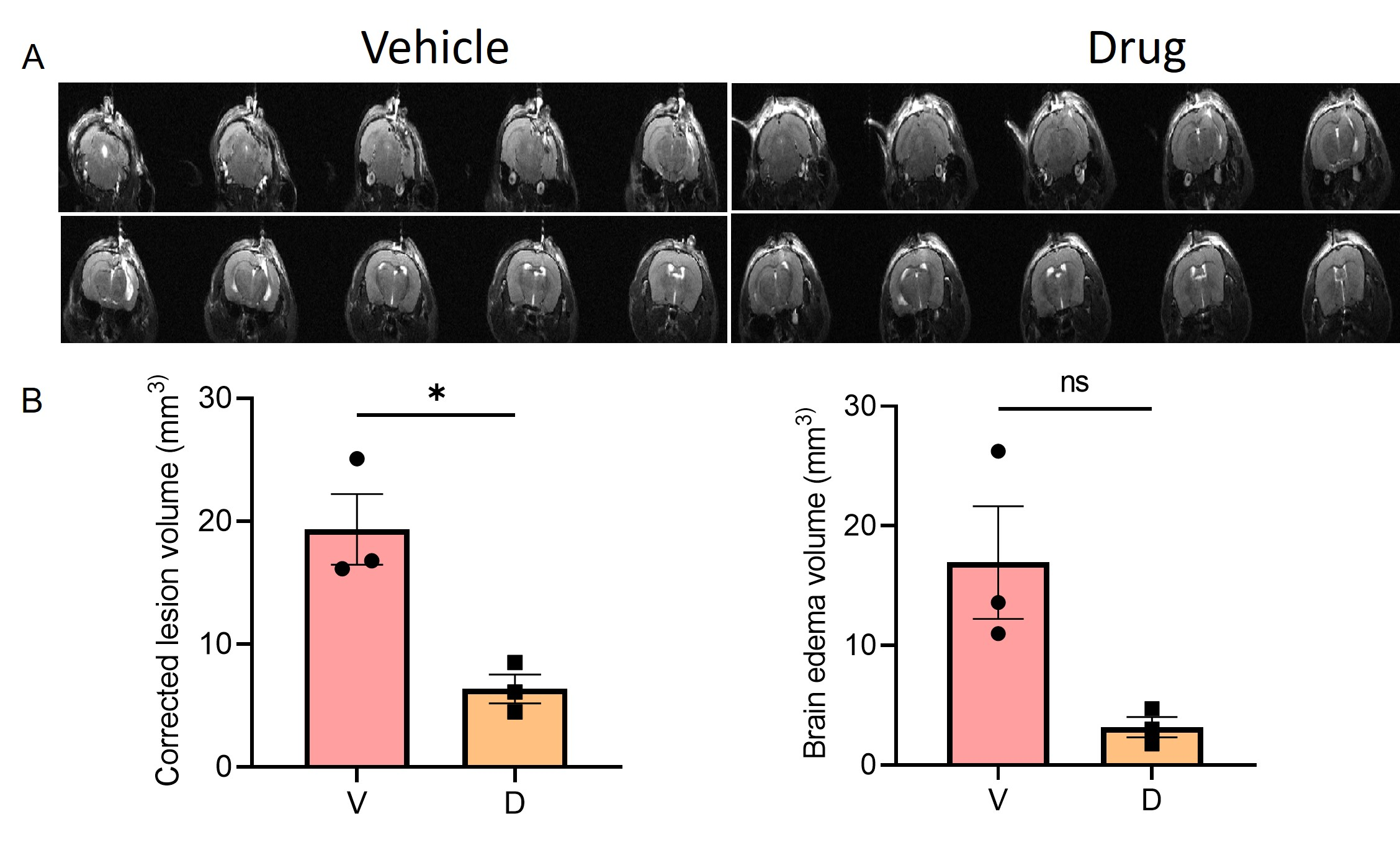

### Suppl fig 6

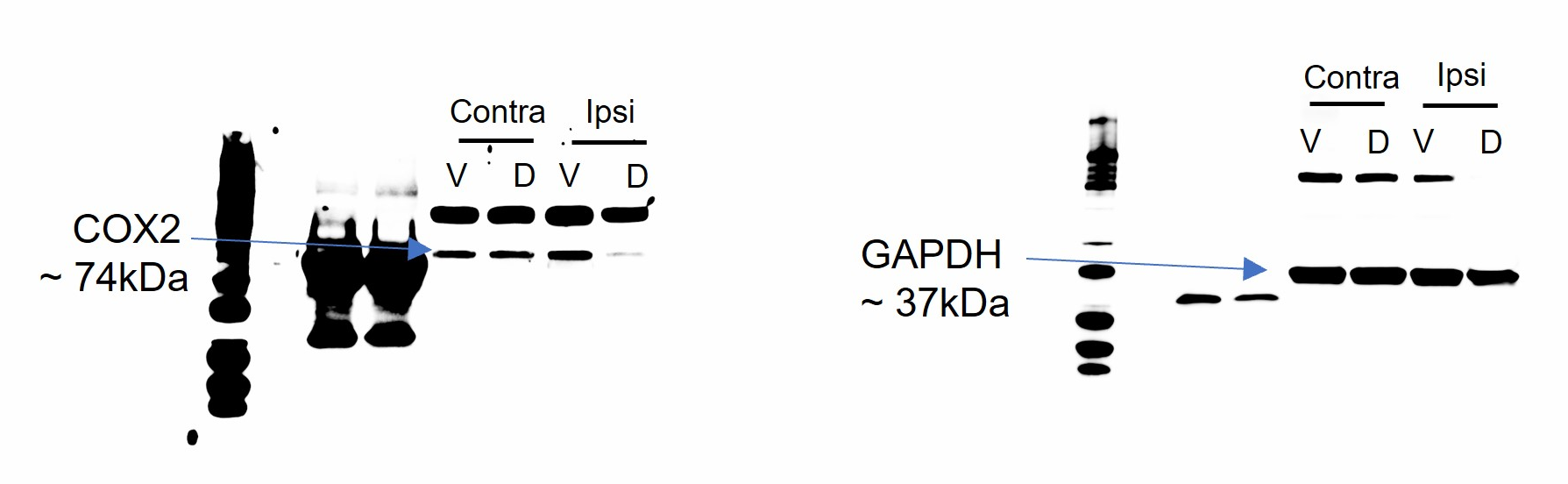

### Suppl fig 7

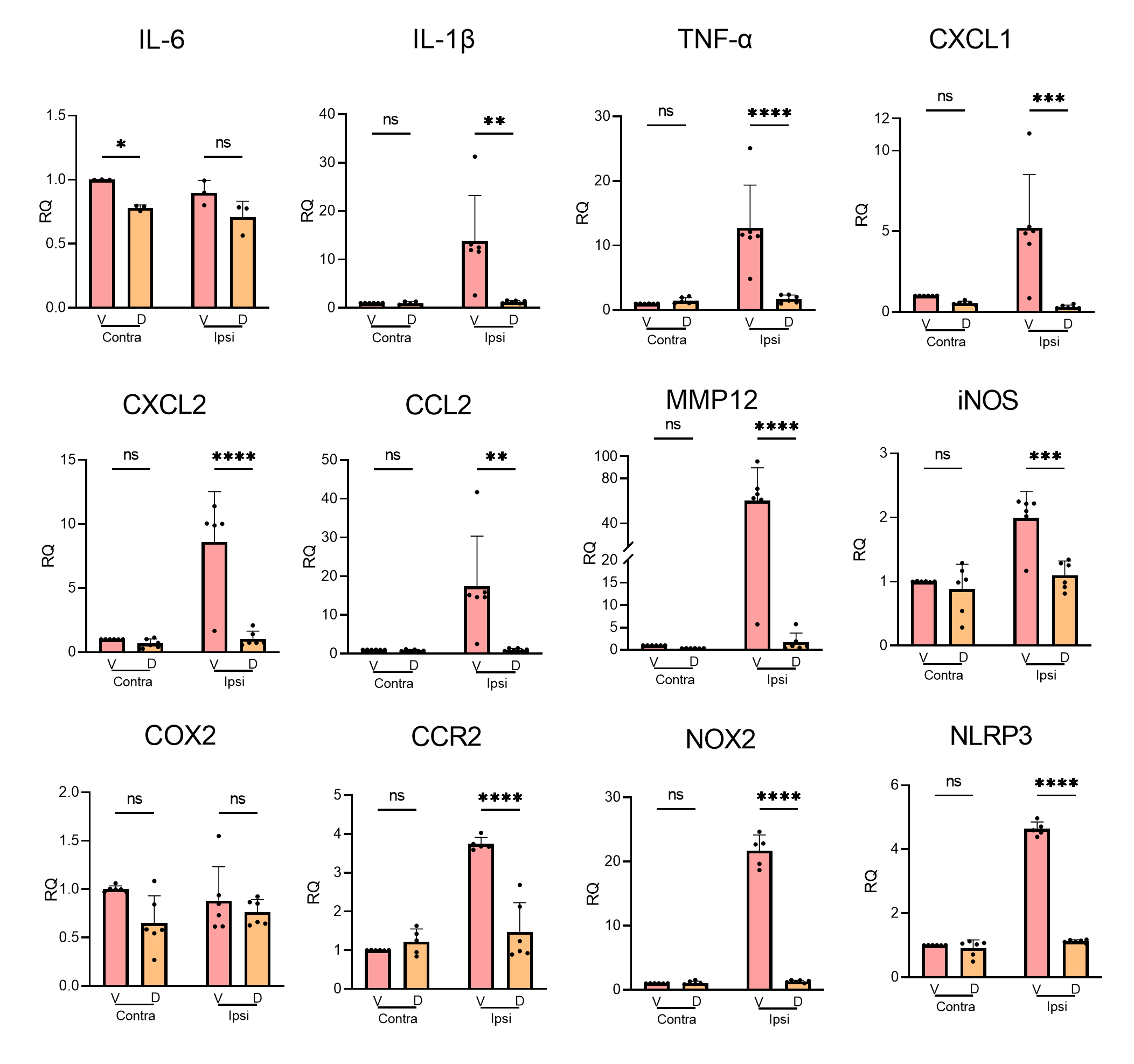

### Suppl fig 8

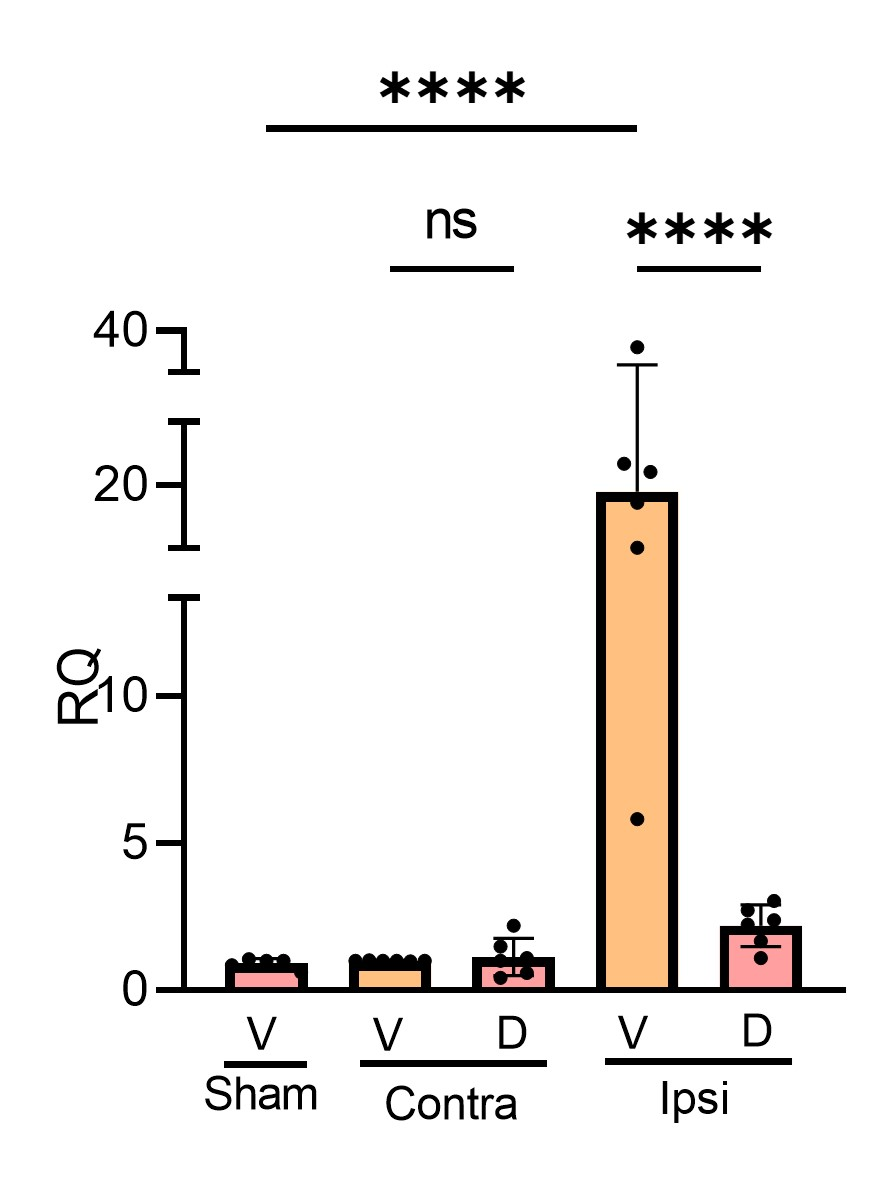
